# CTCF, WAPL and PDS5 proteins control the formation of TADs and loops by cohesin

**DOI:** 10.1101/177444

**Authors:** Gordana Wutz, Csilla Várnai, Kota Nagasaka, David A Cisneros, Roman Stocsits, Wen Tang, Stefan Schoenfelder, Gregor Jessberger, Matthias Muhar, M Julius Hossain, Nike Walther, Birgit Koch, Moritz Kueblbeck, Jan Ellenberg, Johannes Zuber, Peter Fraser, Jan-Michael Peters

**Author notes:** These authors contributed equally to this work (Gordana Wutz, Csilla Várnai, Kota Nagasaka, David A Cisneros).

## Abstract

Mammalian genomes are organized into compartments, topologically-associating domains (TADs) and loops to facilitate gene regulation and other chromosomal functions. Compartments are formed by nucleosomal interactions, but how TADs and loops are generated is unknown. It has been proposed that cohesin forms these structures by extruding loops until it encounters CTCF, but direct evidence for this hypothesis is missing. Here we show that cohesin suppresses compartments but is essential for TADs and loops, that CTCF defines their boundaries, and that WAPL and its PDS5 binding partners control the length of chromatin loops. In the absence of WAPL and PDS5 proteins, cohesin passes CTCF sites with increased frequency, forms extended chromatin loops, accumulates in axial chromosomal positions (vermicelli) and condenses chromosomes to an extent normally only seen in mitosis. These results show that cohesin has an essential genome-wide function in mediating long-range chromatin interactions and support the hypothesis that cohesin creates these by loop extrusion, until it is delayed by CTCF in a manner dependent on PDS5 proteins, or until it is released from DNA by WAPL.

## Introduction

Duplicated DNA molecules become physically connected with each other during DNA replication. This sister chromatid cohesion is essential for bi-orientation of chromosomes on the mitotic or meiotic spindle and thus enables their symmetrical segregation during cell division (Dewar *et al*, 2004). Cohesion is mediated by cohesin complexes (Guacci *et al*, 1997; Michaelis *et al*, 1997; Losada *et al*, 1998) which are thought to perform this function by entrapping both sister DNA molecules inside a ring structure that is formed by the cohesin subunits SMC1, SMC3 and SCC1 (also known as Rad21 and Mcd1) (Haering *et al*, 2008).

Cohesin is present at centromeres and on chromosome arms (reviewed in (Peters *et al*, 2008)). At centromeres, cohesin resists the pulling force of spindle microtubules, a function that is required both for stabilization of microtubule-kinetochore attachments and chromosome bi-orientation. On chromosome arms, however, the precise location of cohesin would not be expected to matter if cohesin’s only function was to mediate cohesion. But contrary to this expectation, cohesin is enriched at thousands of well-defined loci on chromosome arms. In mammalian genomes, ∼90% of these are defined by binding sites for CCCTC binding factor CTCF (Parelho *et al*, 2008; Wendt *et al*, 2008). CTCF is a zinc-finger protein that has been implicated in various aspects of gene regulation, such as insulating gene promoters from distant enhancers (reviewed in (Wendt & Peters, 2009)). Early work revealed that this enhancer blocking activity is required for imprinted gene expression at the *H19-IGF2 locus* (Bell & Felsenfeld, 2000; Hark *et al*, 2000) and indicated that CTCF mediates this function by creating allele specific chromatin loops (Kurukuti *et al*, 2006; Splinter *et al*, 2006). Remarkably, cohesin is also required for CTCF’s enhancer blocking activity at the *H19-IGF2 locus* (Wendt *et al*, 2008) and the chicken *β*-globin locus (Parelho *et al*, 2008), as well as for regulation of other genes (reviewed in (Wendt & Peters, 2009; Dorsett & Merkenschlager, 2013)). Importantly, these requirements have been identified in G1-phase of the cell cycle and in post-mitotic cells, in which cohesin is also present, indicating that they are not an indirect effect of cohesin’s role in sister chromatid cohesion but reflect an independent function of cohesin (Pauli *et al*, 2008; Wendt *et al*, 2008).

Because cohesin is required for CTCF dependent gene regulation events that are thought to be mediated by chromatin looping, and because cohesin is well known to be able to physically connect distinct DNA elements to generate cohesion, it has been proposed that cohesin can also generate or stabilize chromatin loops (Wendt *et al*, 2008; Wendt & Peters, 2009). According to this hypothesis, cohesin would not only be able to connect two sister DNA molecules in *trans* to generate cohesion, but would also be able to connect regions on the same chromatid in *cis*. This hypothesis has been supported by chromatin conformation capture (3C) experiments which indicated that long-range chromosomal *cis*-interactions at the apolipoprotein gene cluster, the *γ-interferon* and *H19-IGF2 loci* (Hadjur *et al*, 2009; Mishiro *et al*, 2009; Nativio *et al*, 2009) as well as enhancer-promoter interactions (Kagey *et al*, 2010) are dependent on cohesin.

Evidence for a role of cohesin in chromatin structure has also come from experiments in which the cohesin associated protein WAPL was depleted. WAPL can release cohesin from DNA (Gandhi *et al*, 2006; Kueng *et al*, 2006; Tedeschi *et al*, 2013), presumably by opening the cohesin ring (Kueng *et al*, 2006; Chan *et al*, 2012; Huis in ’t Veld *et al*, 2014), and WAPL depletion, therefore, increases residence time and amounts of cohesin on DNA (Kueng *et al*, 2006; Tedeschi *et al*, 2013). Remarkably, this causes mild but global compaction of chromatin (Tedeschi *et al*, 2013), indicating that the effects of cohesin on chromatin architecture are widespread and not confined to a few *loci*.

Unexpectedly, WAPL depletion also causes a dramatic intra-nuclear re-localization of cohesin. Cohesin is normally detectable in most regions of interphase chromatin (Losada *et al*, 1998; Sumara *et al*, 2000), but after WAPL depletion accumulates in axial structures called “vermicelli” which are thought to extend from one telomere to the other in interphase chromosome territories (Tedeschi *et al*, 2013). Vermicelli are reminiscent of axial elements in mitotic and meiotic chromosomes. These structures contain condensin and meiosis-specific cohesin complexes, respectively, and are thought to form the base of chromatin loops, into which chromatin fibers are organized during chromosome condensation (Earnshaw & Laemmli, 1983; Saitoh *et al*, 1994; Klein *et al*, 1999; Blat *et al*, 2002; Ono *et al*, 2003; Yeong *et al*, 2003). It has, therefore, been proposed that vermicelli represent cohesin complexes that are located at the base of loops in interphase chromatin, and that vermicelli become detectable only after WAPL depletion because increased residence time and amounts of cohesin lead to the formation or stabilization of more chromatin loops than normally (Tedeschi *et al*, 2013).

Interestingly, both chromatin compaction and accumulation of cohesin axial structures are predicted to occur in WAPL depleted cells if one assumes that cohesin forms chromatin loops *via* a hypothetical loop extrusion process (Fudenberg *et al*, 2016). According to this idea, two distal chromosomal elements would be brought into direct proximity by a loop extrusion factor such as cohesin which would processively extrude a chromatin loop (Riggs, 1990; Nasmyth, 2001; Alipour & Marko, 2012; Gruber, 2014; Sanborn *et al*, 2015; Fudenberg *et al*, 2016). Since WAPL depletion increases cohesin’s residence time on DNA, cohesin would extrude longer loops and cohesin complexes would come into closer proximity than normally, resulting in chromatin compaction and vermicelli formation, respectively. *In silico* modelling of DNA folding has confirmed these predictions (Fudenberg *et al*, 2016).

Importantly, the loop extrusion hypothesis can also explain the “CTCF convergence rule” (Rao *et al*, 2014; de Wit *et al*, 2015; Vietri Rudan *et al*, 2015). This describes the unexpected phenomenon that CTCF binding sites that form the base of chromatin loops are typically oriented towards each other. The DNA sequences with which CTCF can associate are not palindromic (symmetrical), i.e. could be positioned relative towards each other in convergent, tandem or divergent orientations. If CTCF sites formed loops by random association, they would be expected to occur in convergent, tandem and divergent orientations with frequencies of 25%, 50% and 25%, respectively. However, high-resolution genome-wide chromatin conformation capture experiments coupled with DNA sequencing (“Hi-C”) (Lieberman-Aiden *et al*, 2009) have revealed that most chromatin loops contain convergent CTCF sites (Rao *et al*, 2014; de Wit *et al*, 2015; Vietri Rudan *et al*, 2015). This would be expected if these sites were brought into proximity by loop extrusion, but cannot be explained if these sites found each other by random diffusion (Sanborn *et al*, 2015; Fudenberg *et al*, 2016). Based on these observations and considerations it has been proposed that CTCF sites function as directional boundary elements during loop extrusion (Sanborn *et al*, 2015; Fudenberg *et al*, 2016). However, for cohesin to be able to form extended loops in WAPL depleted cells, as proposed by Fudenberg et al, (Fudenberg *et al*, 2016), these boundaries would have to be permeable to some extent.

The notion that CTCF sites can function as boundary elements for loop extrusion is consistent with the observation that cohesin and CTCF co-localize (Parelho *et al*, 2008; Wendt *et al*, 2008), and that many of these sites either form the base of chromatin loops (Rao *et al*, 2014), or are enriched at the boundaries of more complex chromatin structures, called topologically associating domains (TADs) (Dixon *et al*, 2012; Nora *et al*, 2012). These are covering chromosomal regions, typically up to 1 Mb in length, in which long-range chromosomal *cis*-interactions occur with increased frequencies. At the level of individual cells, TADs may represent loops that are in the process of being extruded to various degrees. Consistent with this possibility, single-nucleus Hi-C experiments indicate that TADs emerge from averaging interactions in large cell populations (Flyamer *et al*, 2017). Importantly, genome editing experiments in which CTCF sites were deleted or inverted indicate that these are indeed required for the formation of TAD boundaries (Nora *et al*, 2012; de Wit *et al*, 2015; Narendra *et al*, 2015; Sanborn *et al*, 2015).

The hypothesis that cohesin contributes to the formation of TADs and loops by mediating loop extrusion is also consistent with the observation that cohesin can move along DNA, both *in vitro* (Davidson *et al*, 2016; Kanke *et al*, 2016; Stigler *et al*, 2016) and *in vivo* (Lengronne 2004; Busslinger *et al*, 2017) and is constrained during these movements by CTCF (Davidson *et al*, 2016; Busslinger *et al*, 2017). In contrast, cohesin and CTCF are not thought to contribute to chromatin organization at levels above TADs and loops (i.e. covering larger genomic regions), at which Hi-C experiments have identified higher-order structures called A and B compartments (Lieberman-Aiden *et al*, 2009). Both of these exist in different sub-types (Rao *et al*, 2014) and are thought to represent euchromatic and heterochromatic regions, respectively.

However, even though the above-mentioned observations indicate that cohesin and CTCF may have important roles in the formation of TADs and loops, previous Hi-C experiments found that inactivation of cohesin or CTCF had only modest effects on chromatin organization (Seitan *et al*, 2013; Sofueva *et al*, 2013; Zuin *et al*, 2014). It is, therefore, incompletely understood whether cohesin and CTCF have important roles in chromatin organization, and if so, whether these are mediated by loop extrusion.

We have, therefore, further analyzed the roles of cohesin and CTCF in chromatin organization by using microscopic imaging and Hi-C experiments. For this purpose, we have either inactivated cohesin or CTCF by conditional proteolysis (Nishimura *et al*, 2009), or stabilized cohesin on DNA by depleting WAPL and its binding partners PDS5A and PDS5B (Kueng *et al*, 2006). Our experiments indicate that cohesin is required for the formation and/or maintenance of TADs and loops but suppresses compartments, that WAPL and PDS5 proteins antagonize these functions, and that CTCF is required for the formation of “sharp” boundaries between TADs and the formation of loops by defined loop anchors. We also show that co-depletion of WAPL, PDS5A and PDS5B enables cohesin to condense chromosomes to an extent that is normally only seen in early mitosis by creating long-range chromosomal *cis*-interactions that are reminiscent of the ones observed in mitotic chromosomes (Naumova *et al*, 2013; Nagano *et al*, 2017). These observations are consistent with the loop extrusion hypothesis. Unexpectedly, however, our results indicate that PDS5 proteins, which are thought to cooperate with WAPL in releasing cohesin from DNA (Rowland *et al*, 2009; Shintomi & Hirano, 2009; Ouyang *et al*, 2016), have a function in chromatin loop formation that is distinct from the role of WAPL and that may be required for CTCF’s ability to function as a boundary element for loop extrusion.

Cohesin, CTCF and WAPL, but not PDS5A and PDS5B, have also been inactivated in other recent Hi-C studies which during preparation of this manuscript were deposited on *bioRxiv* (Schwarzer *et al*, 2016; Kubo *et al*, 2017; Rao *et al*, 2017) or published in peer reviewed journals (Haarhuis *et al*, 2017; Nora *et al*, 2017). We compare and contrast the results of these as well as the earlier studies (Seitan *et al*, 2013; Sofueva *et al*, 2013; Zuin *et al*, 2014) with ours in the Discussion.

## Results

### Acute cohesin inactivation by auxin mediated degradation

To analyze cohesin’s role in chromatin structure, we modified all SCC1 alleles in Hela cells by CRISPR-Cas9 mediated genome editing to encode proteins in which SCC1 is C-terminally fused to green fluorescent protein (GFP) and an auxin inducible degron (AID; Fig 1A). To enable auxin dependent recognition of SCC1-GFP-AID by SCF ubiquitin ligases, we stably expressed the *Oryza sativa* F-box transport inhibitor response-1 auxin receptor protein (Tir1) in these cells. In immunoblotting experiments, only SCC1-GFP-AID but no untagged SCC1 could be detected, confirming that all SCC1 alleles had been modified (Fig 1B). These experiments also revealed that GFP-AID tagging reduced SCC1 levels (Fig 1B), consistent with the finding that C-terminal tagging compromises SCC1 function in mice (Tachibana-Konwalski *et al*, 2010). Cells expressing only SCC1-GFP-AID nevertheless proliferated at similar rates as wild type cells, indicating that cohesin complexes containing this fusion protein can perform their essential cellular functions and are present in copy numbers sufficient for this.

**Figure 1.**
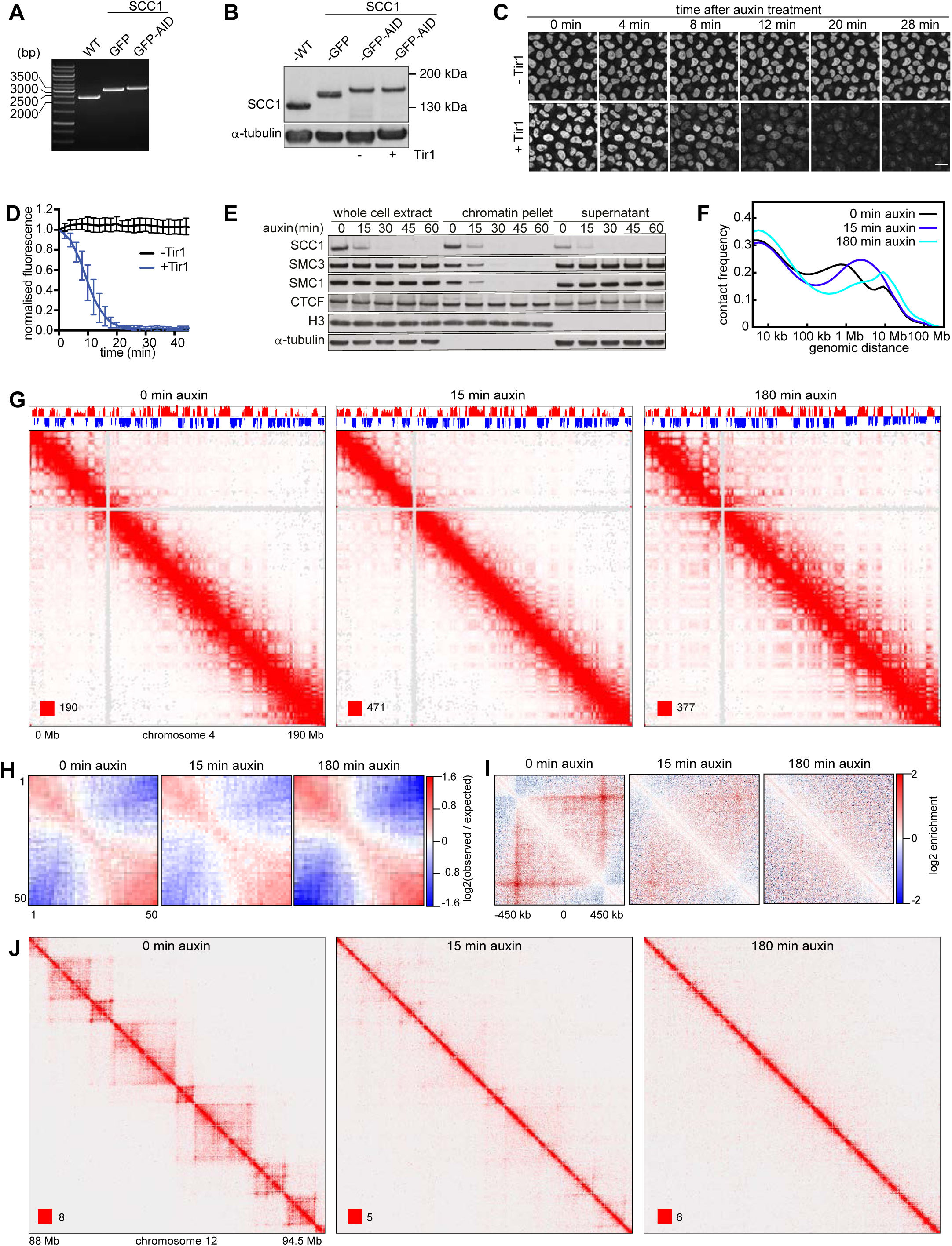
Chromatin organisation changes upon auxin induced SCC1 degradation. A. Genotype analysis of parental HeLa cells (WT), homozygous SCC1-GFP cells (GFP) and homozygous SCC1-GFP-AID cells (GFP-AID). Genomic PCR products were generated with the primers that are designed external to the homology arm which was used for inserting GFP or GFP-AID encoding sequences downstream of the SCC1 gene. This resulted in fusion proteins with GFP or GFP-AID tags C-terminal to the SCC1 gene. B. Immunoblotting analysis of whole cell extracts from the cell lines characterized in WT, SCC1-GFP, SCC1-GFP-AID (-) cells (i.e. not expressing Tir1) and SCC1-GFP-AID cells expressing Tir1 (+). a-tubulin: loading control. C. Time course live cell imaging of SCC1-GFP-AID cells after auxin treatment. SCC1-GFP-AID cells with (+) or without (-) Tir1 were imaged after addition of auxin. Scale bar indicates 20 μm. D. Quantification of nuclear GFP signal over time after auxin addition to SCC1-GFP-AID cells with (+) or without (-) Tir1. Error bar depicts standard deviations of the means. E. Chromatin fractionation and immunoblot analysis of auxin-treated SCC1-GFP-AID cells expressing Tir1. At the indicated time points after auxin addition, whole cell extracts, the chromatin pellet fraction and the supernatant fraction were analyzed by immunoblotting, using antibodies against the proteins indicated on the left. F. Intra-chromosomal contact frequency distribution as a function of genomic distance, at 0 (black), 15 (blue) and 180 minutes (cyan) after auxin addition to SCC1-GFP-AID cells expressing Tir1. G. Coverage-corrected Hi-C contact matrices of chromosome 4, at 0 (left), 15 (centre) and 180 minutes (right) after auxin addition to SCC1-GFP-AID cells expressing Tir1. The corresponding compartment signal tracks at 250kb bin resolution are shown above the matrices. The matrices were plotted using Juicebox. H. Long-range (>2Mb) intra-chromosomal contact enrichment between bins with varying compartment signal strength from most B-like (1) to most A-like (50). I. Average contact enrichment around loops after auxin addition to SCC1-GFP-AID cells expressing Tir1, for the 82 x 600 kb long loops identified in G1 control HeLa cells. The matrices are centered (0) around the halfway point of the loop anchor coordinates. J. For the same conditions as in (G-I), coverage-corrected Hi-C contact matrices in the 88-94.5Mb region of chromosome 12, plotted using Juicebox.

Auxin addition reduced GFP fluorescence intensity to “background” levels in 20 min in cells expressing Tir1, but not in cells lacking Tir1 (Figs 1C and D). Immunoblotting experiments confirmed that loss of fluorescence intensity was caused by SCC1-GFP-AID degradation and showed that it resulted in concomitant release of the cohesin subunits SMC1 and SMC3 from chromatin, whereas CTCF and the cohesin loading complex NIPBL-MAU2 remained chromatin associated (Fig 1E and EV1A).

### Cohesin inactivation strengthens compartments but destroys TADs and loops

To analyze the effects of cohesin inactivation on chromatin structure, we synchronized SCC1-GFP-AID cells in G1-phase by double thymidine arrest-release, treated them for 0, 15, or 180 minutes with auxin, and measured chromatin interactions genome wide by Hi-C (Fig 1F-J). In a separate experiment, we analyzed the same cells after 0 and 120 min of auxin treatment (Appendix Fig S1; all Hi-C libraries analyzed in this study and their properties are listed in Appendix Table S1). All genome-wide interaction matrices were analyzed for the presence of 5 types of patterns which have been observed in Hi-C data from mammalian genomes: *cis-trans* interaction ratios and distance-dependent interaction frequencies at the genomic level, and compartments, TADs and point interactions (“loops”) at specific genomic positions (Lajoie *et al*, 2015). Importantly, none of these patterns differed significantly between Hi-C matrices obtained from cells collected at the 0-min time points of the two experiments, indicating that changes observed in auxin treated cells were caused by cohesin degradation and not inter-experimental variation.

*Cis-trans* interaction ratios were high in all cases, with inter-chromosomal interactions only representing 5-15%, indicating that all Hi-C libraries were of high quality. However, *trans* interactions were consistently higher in cohesin depleted cells (Appendix Table S1). Contact frequencies were affected differently by cohesin inactivation, depending on genomic distances between contacting loci. Cohesin depletion gradually decreased contact frequencies at 100 kb-1 Mb distances, while the frequency of long-range (>10 Mb) interactions increased (Fig 1F). Cohesin inactivation might therefore enable more contacts with distant parts of chromosomes and even neighbouring chromosomes, possibly because chromatin structure becomes more flexible under these conditions. For reasons explained in the next paragraph, we do not believe that these changes are caused by technical artefacts, such as increased “noise” in the Hi-C libraries.

Cohesin degradation also induced major changes at the level of compartments, TADs and loops. In cells treated for 120 or 180 min with auxin, A and B compartments could be detected more clearly than in control cells (Appendix Table 1), as coverage-corrected maps of single chromosomes showed increased interaction frequencies further away from the diagonal of the Hi-C matrices, resulting in enhanced “plaid” or “checkerboard” patterns (Fig 1G). Aggregate analysis of 50 compartment categories ranging from strong B to strong A compartments genome-wide confirmed this, showing increasing contact enrichment between similar compartment categories and a decreasing contact enrichment between dissimilar (e.g. strong A and strong B) compartment bins in both long *cis* (>2 Mb, Fig 1H) and *trans* interactions. The strengthened compartmentalization suggests that the increase in long-range contact frequencies described above was not due to “noise” in the Hi-C libraries, since more long-range random contacts would result in less contact specificity and a weakening in compartmentalization strength.

Whereas compartments became more apparent in cohesin depleted cells, TADs were greatly reduced (Fig 1J). After auxin addition, there was a gradual decrease in TAD detectability based on directionality indices (Dixon *et al*, 2012), with the genome coverage of identified TADs decreasing from 56% in control cells to 52% and 44% after 15 and 180 min of auxin treatment, respectively. The insulation score calculated for TAD borders declined concomitant with the disappearance of TADs (Fig EV1B). Individual loops also disappeared gradually following cohesin degradation until they were barely detectable after 180 min; in this experiment “juicer-tool” (Rao *et al*, 2014) identified 5,740 loops in control cells, but only 204 (3.6%) after 180 min of auxin treatment (Fig EV1C; aggregate analyses of loops 600 kb in length are shown in Fig 1I and of 0.75-6 Mb loops in Fig EV1D). In this context, it is important to note that the number of loops that can be called depends on the number of replicate Hi-C libraries sequenced, with more replicate numbers yielding higher signal-to-noise ratios and thus higher loop numbers. In each given experiment of this study we, therefore, compared data from identical numbers of replicates. However, because different replicate numbers were sequenced in different experiments, the absolute number of loops identified varies between experiments (see below).

These results indicate that local genome organization into TADs and loops depends critically on cohesin and changes rapidly after its inactivation, whereas cohesin suppresses the formation of A and B compartments or their detectability by Hi-C. Interestingly, after 15 min auxin treatment, 96% of loops but only 53% of TADs had disappeared, whereas enhanced compartmentalization was only observed after 180 min of auxin treatment. Loops are therefore particularly sensitive to cohesin inactivation, whereas changes in TADs occur more slowly, and changes in compartments on an even longer time scale.

For comparison, we also analyzed HeLa cells in which SCC1 was not tagged (Appendix Fig S2). Unexpectedly, contact frequencies at 20 kb-1 Mb distances were higher in these than in SCC1-AID-GFP cells, while the frequency of long-range (>2 Mb) interactions was lower (Appendix Fig S2A). The boundary insulation score and the number of loops that could be called were also higher in unmodified than in SCC1-AID-GFP expressing cells, as if chromatin was more highly organized in the former than the latter cells (Appendix Figs S2B and C). We suspect that these differences are caused by the reduced expression levels of SCC1-AID-GFP (see Fig 1B), and/or by functional defects caused by SCC1 tagging, such as reduced rate or processivity of loop extrusion.

### Acute CTCF inactivation by auxin mediated degradation

To analyze CTCF’s role in chromatin structure, we generated Hela cells in which all CTCF alleles were modified to encode CTCF-GFP-AID (Fig 2A). Immunoblotting revealed that Tir1 expression reduced CTCF-GFP-AID levels even in the absence of auxin (Fig 2B), implying that Tir1 targets some CTCF-GFP-AID for degradation on its own. However, also these cells proliferated at similar rates as wild type cells, indicating that the functionality and copy number of CTCF-GFP-AID are sufficient for cell viability and proliferation.

**Figure 2.**
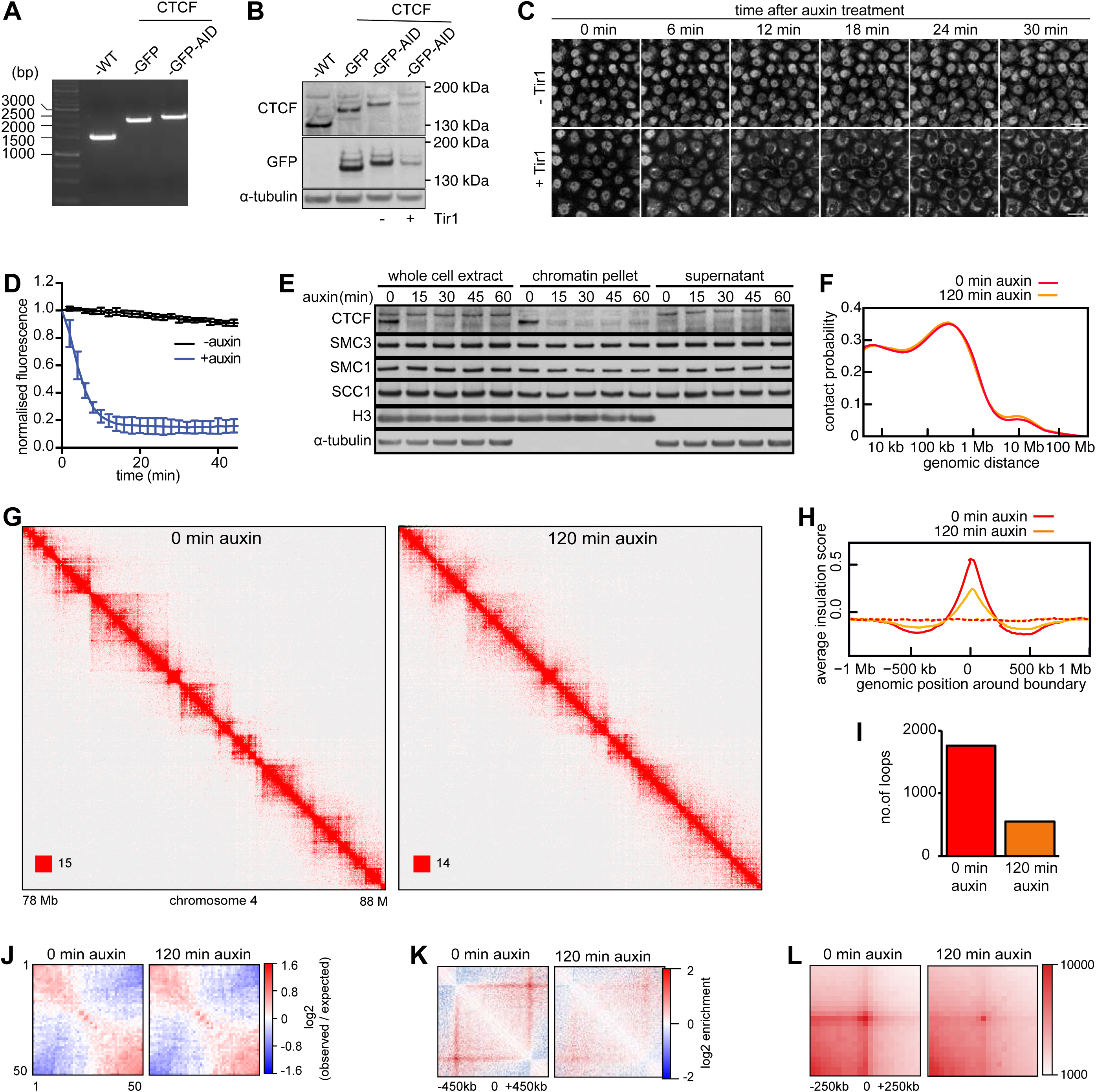
Chromatin organisation changes upon auxin-induced CTCF degradation. A. Genotype analysis of parental HeLa cells (WT), homozygous CTCF-GFP cells (GFP) and homozygous CTCF-GFP-AID cells (GFP-AID). Genomic PCR products were generated with the primers that are designed external to the homology arm which was used for inserting GFP or GFP-AID encoding sequences downstream of the CTCF gene. This resulted in fusion proteins with GFP or GFP-AID tags C-terminal to the CTCF gene. B. Immunoblotting analysis of whole cell extracts from the cell lines characterized in WT, CTCF-GFP, CTCF-GFP-AID (-) cells (i.e. not expressing Tir1) and CTCF-GFP-AID cells expressing Tir1 (+). a-tubulin: loading control. C. Time course live cell imaging of CTCF-GFP-AID cells after auxin treatment. CTCF-GFP-AID cells with (+) or without (-) Tir1 were imaged after addition of auxin. Scale bar indicates 20 μm. D. Quantification of nuclear GFP signal over time after auxin addition to CTCF-GFP-AID cells with (+) or without (-) Tir1. Error bar depicts standard deviations of the means. E. Chromatin fractionation and immunoblot analysis of auxin-treated CTCF-GFP-AID cells expressing Tir1. At the indicated time points after auxin addition, whole cell extracts, the chromatin pellet fraction and the supernatant fraction were analyzed by immunoblotting, using antibodies against the proteins indicated on the left. F. Intra-chromosomal contact frequency distribution as a function of genomic distance, 0 (orange) and 120 minutes (red) after auxin addition to CTCF-GFP-AID cells expressing Tir1. G. Coverage-corrected Hi-C contact matrices of chromosome 4 (78-88Mb), 0 (left) and 120 minutes (right) after auxin addition to CTCF-GFP-AID cells expressing Tir1. The matrices were plotted using Juicebox. H. Average insulation score around TAD boundaries identified in G1 control cells, for samples at 0 (red) and 120 minutes (yellow) after auxin addition to CTCF-GFP-AID cells expressing Tir1. I. Number of loops identified by HICCUPS, in the same conditions as (F-H). J. Long-range (>2Mb) intra-chromosomal contact enrichment between bins with varying compartment signal strength from most B-like (1) to most A-like (50), in the same conditions as (F). K. Average contact enrichment around loops after auxin addition to CTCF-GFP-AID cells expressing Tir1, for the 82 x 600 kb long loops identified by HICCUPS in G1 control HeLa cells. The matrices are centered (0) around the halfway point of the loop anchor coordinates. L. Total contact counts around loops after auxin addition, for all 750kb-6Mb long loops identified by HICCUPS in G1 control. The vertical and horizontal axes of the matrices were centered around the upstream and the downstream loop anchors, respectively.

Auxin addition reduced the fluorescence intensity of CTCF-GFP-AID to “background” levels in 15 min (Figs 2C and D). GFP signals were low in these cells, consistent with the low CTCF-GFP-AID levels observed by immunoblotting. CTCF-GFP-AID could thus only be detected under imaging conditions which also revealed cytoplasmic fluorescence. However, this cytoplasmic signal did not disappear following auxin treatment, indicating that it was caused by auto-fluorescent molecules other than GFP. Immunoblotting confirmed that auxin induces rapid degradation of most CTCF-GFP-AID (Fig 2E).

### CTCF inactivation reduces the insulation between TADs

To analyze the effects of CTCF inactivation on chromatin structure we synchronized CTCF-GFP-AID cells in G1-phase, treated them for 0 and 120 minutes with auxin, and measured chromatin interactions genome wide by Hi-C (Fig 2F-L). CTCF degradation did not change contact frequencies at any contact distances (Fig 2F), including inter-chromosomal contacts (Appendix Table S1). The long-range *cis* and *trans* chromosomal compartmentalization strength was also unaffected (Fig 2J and Appendix Table S1). However, CTCF inactivation had a pronounced effect on TAD boundaries, which appeared fuzzier and less well defined than in control cells (Fig 2G). This was reflected by a strong decrease in the TAD insulation score (Fig 2H) and resulted in a reduction in TAD number as identified by directionality indices (from 58.4% to 50.8%). Likewise, most loops disappeared in cells depleted of CTCF; “juicer-tool” identified 1,763 loops before auxin addition, but only 546 loops 120 min after auxin addition (Figs 2I). This number of loops (1,763) is much smaller than the number identified in unmodified Hela cells (5,740; Appendix Fig S2C), in which also the TAD insulation score is higher (Appendix Fig S2B). As for SCC1-GFP-AID cells, we, therefore, suspect that CTCF-GFP-AID cells are hypo-morphic with respect to CTCF function, either because of their reduced CTCF levels or because of functional interference from the GFP-AID tag.

Together, these results indicate that CTCF has a distance-independent role in chromatin organization, is not essential for compartmentalization, but is required for the formation and/or maintenance of TAD borders and loops. These observations imply that CTCF, unlike cohesin, is not required for long-range chromatin interactions *per se*, but has an important role in defining their precise position, consistent with the proposed role of CTCF as a boundary for loop extrusion (Sanborn *et al*, 2015; Fudenberg *et al*, 2016).

### PDS5 proteins cooperate with WAPL in controlling cohesin localization and chromatin structure

Our results so far indicated that cohesin is required for the formation and/or maintenance of TADs and loops, but suppresses compartmentalization, whereas CTCF is required for the formation of loops and distinct borders between TADs. To test these hypotheses further, we analyzed if and how genome structure is altered in WAPL depleted cells, in which cohesin cannot be properly released from DNA (Tedeschi *et al*, 2013). We were also interested in understanding how WAPL depletion causes mild chromatin compaction and accumulation of cohesin in axial structures (“vermicelli”), and we wanted to know if and how WAPL’s binding partners PDS5A and PDS5B contribute to these processes.

To address these questions, we first depleted WAPL, PDS5A and PDS5B (in the figures abbreviated as W, A and B, respectively) either singly or in combination, and analyzed cohesin distribution and chromatin morphology in fixed cells. For this purpose, we used Hela cells in which all SCC1 alleles were modified to encode SCC1-GFP (Figs 1A and B) (Davidson *et al*, 2016), enabling us to analyze endogenous functional cohesin. Because of the large number of possible combinations in which WAPL, PDS5A and PDS5B could be depleted, we used RNA interference (RNAi) instead of AID tagging in these experiments. Immunoblot analyses indicated that RNAi enabled depletion efficiencies of >88% for PDS5B and >94% for PDS5A and WAPL (Fig 3A).

**Figure 3.**
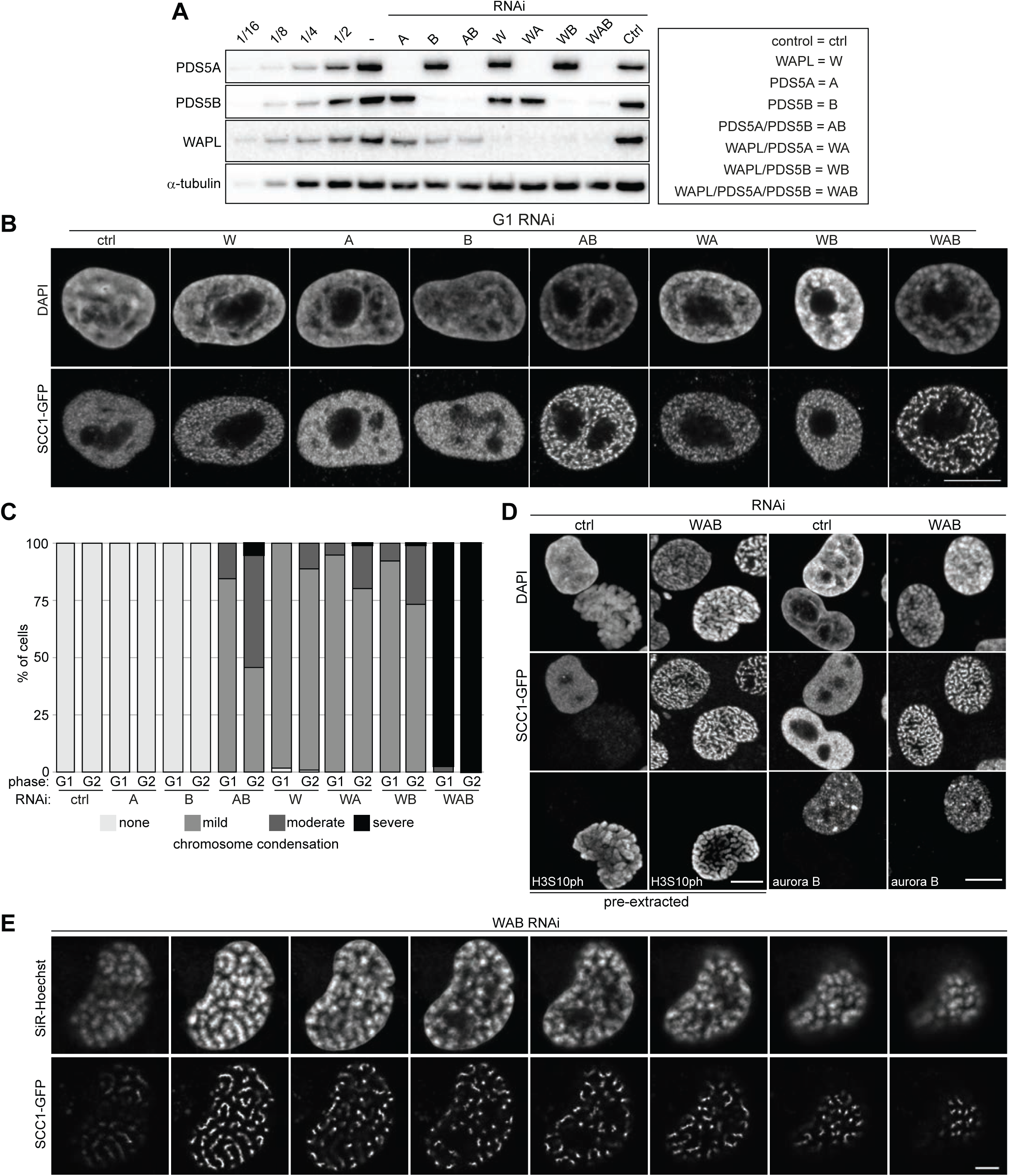
Depletion of PDS5A/B induces chromatin compaction and vermicelli formation. A. Immunoblotting analysis of SCC1-GFP cells treated with indicated single siRNAs or their combinations. B. Representative immunofluorescence images of SCC1-GFP cells shown in (A) stained for GFP at G1 phase. DNA was counterstained with DAPI. Scale bar shows 10 μm. C. Phenotypic classification of chromatin compaction in G1 and G2 phase in (B) and Appendix Fig S3, n > 100 cells per condition. D. Immunofluorescence microscopy of SCC1-GFP cells depleted for WAB or control-depleted (ctrl). Fixed cells with or without pre-extraction were stained for GFP and DAPI in combination with either phospho-H3 Ser10 (H3S10ph) or Aurora B staining. Scale bar shows 10 μm. E. Live cell imaging of SCC1-GFP cells depleted for WAB. Individual confocal sections from Appendix Fig S3D are shown, using 0.4 μm confocal distance between original sections. DNA was counter stained with SiR-DNA. Scale bar shows 5 μm.

Immunofluorescence microscopy revealed that WAPL depletion caused accumulation of SCC1 in vermicelli, as previously seen in mouse embryonic fibroblasts (MEFs) (Tedeschi *et al*, 2013; Busslinger *et al*, 2017). In contrast, depletion of PDS5A or PDS5B changed cohesin localization a little, and co-depletion of WAPL with either PDS5A or PDS5B did not enhance the vermicelli phenotype seen after depletion of WAPL alone. However, co-depletion of PDS5A and PDS5B caused accumulation of cohesin in vermicelli (Fig 3B), as recently also reported by (Ouyang *et al*, 2016). Vermicelli in cells depleted of both PDS5 proteins were more pronounced than after WAPL depletion, and simultaneous depletion of all three proteins enhanced this phenotype even further (Fig 3B).

Staining of DNA with 4’,6-diamidino-2-phenylindole (DAPI) revealed that WAPL depletion and co-depletion of PDS5A and PDS5B caused mild chromatin compaction, as seen in WAPL depleted MEFs (Tedeschi *et al*, 2013). However, depletion of all three proteins caused strong condensation, resulting in the separation of chromatin into individual chromosomes (Figs 3B and C; Appendix Fig S3A). These were in many cases as condensed as chromosomes in mitotic prophase, even though cells depleted of WAPL, PDS5A and PDS5B remained in interphase, as judged by the absence of histone H3 phosphorylated on serine 10 (H3S10ph; Fig 3D). Instead, co-staining with aurora B antibodies revealed that the cohesin localization and chromatin compaction phenotypes were present in both G1-phase (aurora B negative) and G2-phase (aurora B positive; Fig 3D), with slightly higher penetrance in G2 (Fig 3C and Appendix Fig S3B). Similar phenotypes were observed by live cell imaging, ruling out fixation artefacts (Appendix Fig S3C). Confocal live cell imaging showed that condensed chromosomes tended to be localized near the nuclear periphery (Fig 3E and Appendix Fig S3D), as are chromosomes in prophase.

These results indicate that, like WAPL, PDS5 proteins control cohesin localization and chromatin compaction, that the functions of PDS5A and PDS5B in these processes are redundant, and that depletion of all three proteins causes compaction of chromatin to a remarkably strong extent, as is normally only seen in early mitosis.

### Co-depletion of PDS5 proteins increases the residence time and amounts of cohesin on chromatin

WAPL depletion increases the residence time and amounts of cohesin on interphase chromatin (Kueng *et al*, 2006; Tedeschi *et al*, 2013). We therefore analyzed if depletion of PDS5 proteins has similar effects. Consistent with a recent report (Ouyang *et al*, 2016), inverse fluorescence recovery after photobleaching (iFRAP) experiments, performed in cells synchronized in G1-phase, showed that co-depletion of PDS5A and PDS5B increased the chromatin residence time of cohesin significantly to t1/2 = 12.2 min, compared to 4.8 min in control cells (Figs 4A and B). However, co-depletion of PDS5 proteins stabilized cohesin less on chromatin than depletion of WAPL (t1/2 = 28.9 min) or depletion of all three proteins (t1/2 could not be measured in these cells due to the slow recovery of cohesin; Fig 4C). Despite these differences, co-depletion of PDS5A and PDS5B caused accumulation of cohesin on chromatin to a similar extent as was seen after WAPL depletion, both in quantitative IFM experiments (Figs 4D-F), and in chromatin fractionation experiments in which samples were analyzed by immunoblotting (Fig 4G). In both assays, the increase of cohesin on chromatin was small, presumably because approximately 50% of cohesin is already bound to chromatin in control cells (Gerlich *et al*, 2006). Inactivation of cohesin release can therefore maximally lead to a twofold increase of cohesin on chromatin. This is difficult to detect by immunoblotting (compare lanes 7 and 10-12 in Fig 4G) but can be inferred from the loss of cohesin from supernatant fractions (compare lanes 13 and 16-18).

**Figure 4.**
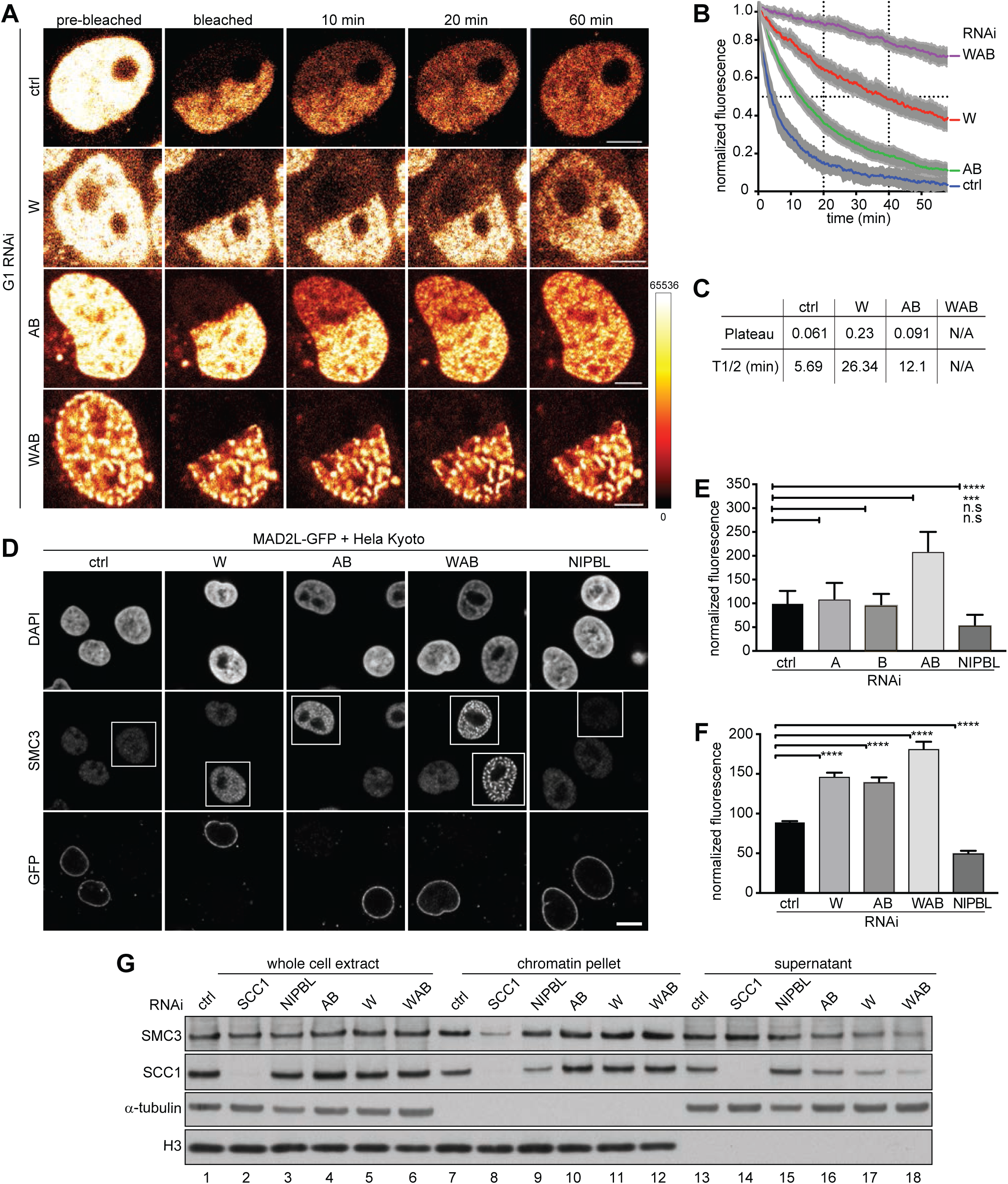
RNAi depletion of PDS5A/B increases the amount and residence time of chromatin-bound cohesin. A. Representative images of an inverse fluorescence recovery after photobleaching (iFRAP) experiment using SCC1-GFP cell lines in G1 phase that were control-depleted (ctrl) or depleted for WAPL (W), PDS5A/B (AB), WAPL/PDS5A/B (WAB). Fluorescence intensities are false coloured as indicated. Scale bar indicates 5 μm. B. Normalised signal intensities after photobleaching, for the iFRAP experiment shown in (A). Error bars denote standard error of the mean (s.e.m), n > 8 cells per condition. C. Quantification of the plateau and half-life of recovery for the curves in (B). D. Immunofluorescence staining of chromatin-bound SMC3 in HeLa cells. HeLa cells were control-depleted (ctrl) or depleted for WAPL (W), PDS5A/B (AB), WAPL/PDS5A/B (WAB) or NIPBL, mixed and seeded with HeLa cells expressing MAD2L-GFP. These cells were pre-extracted prior to fixation and stained for DAPI, SMC3 and GFP. Scale bar indicates 10 μm. E. Quantification of SMC3 fluorescence intensities obtained in the experiments shown in (D). SMC3 fluorescence intensities were normalised to those of neighbouring RNAi-untreated cells marked by MAD2L-GFP cells. RNAi treated cells are entangled. The asterisk denotes a significant difference according to a Dunn’s-corrected Kruskal-Wallis test (n>90, ****= p value ≤ 0.0001). F. Same as panel (E) except for using PDS5A-GFP cell lines as RNAi-untreated cells. The asterisk denotes a significant difference according to a Dunn’s-corrected Kruskal-Wallis test (n>30, ****= p value ≤ 0.0001, ***= p value ≤ 0.001, N.S.=p value>0.05). G. Immunoblotting analysis of the whole cell extract, the chromatin pellet fraction and the supernatant fraction from HeLa cells treated with siRNAs indicated on the left.

These results indicate that both PDS5 proteins cooperate with WAPL in releasing cohesin from chromatin and that WAPL has a more important role in this process, as its depletion increases cohesin’s residence time much more than depletion of PDS5 proteins (Fig 4B). However, the precise contribution of the different proteins is difficult to assess from RNAi experiments because these tend to cause hypomorphic conditions and not “null” phenotypes.

### No evidence that condensin complexes contribute to chromosome condensation in cells depleted of WAPL and PDS5 proteins

Because simultaneous depletion of WAPL and PDS5 proteins causes mitotic-like condensation of chromosomes we analyzed the localization and function of condensin complexes, which have been implicated in mitotic chromosome condensation. For this purpose, we used Hela Kyoto cells in which all alleles of either CAP-D2 or CAP-D3 were modified to encode C-terminal mEGFP fusion proteins, enabling the visualization of endogenous condensin I and condensin II complexes, respectively (Appendix Fig S4A). Whereas co-depletion of WAPL and PDS5 proteins caused the re-localization of cohesin into vermicelli described above, no change in condensin localization could be detected, with condensin I remaining cytoplasmic and condensin II remaining evenly distributed in the nucleus (Appendix Fig S4B). Furthermore, co-depletion of SMC2, a subunit of both condensin I and II (Ono *et al*, 2003; Yeong *et al*, 2003), together with WAPL, PDS5A and PDS5B did not prevent chromosome condensation and vermicelli formation (Appendix Figs S4C and D). These results do not formally exclude a role for condensins because they may not have been fully depleted by RNAi, but they do imply that cohesin is the major cause of chromosome condensation in cells depleted of WAPL, PDS5A and PDS5B.

### Depletion of WAPL, PDS5A and PDS5B weakens compartments but increases the size of TADs

To understand how depletion of WAPL and PDS5 proteins causes vermicelli formation and chromatin compaction we performed Hi-C experiments. For this purpose, we depleted WAPL alone, or PDS5A and PDS5B together, or all three proteins in cells synchronized in G1-phase and compared them to cells transfected with control siRNAs. For comparison, we also analyzed cells not transfected with siRNAs which had been synchronized in S-phase, G2-phase or prometaphase (see below). All Hi-C experiments were performed as independent biological replicates. Cell samples from the same populations that were used in Hi-C experiments were analyzed for DNA content by fluorescence activated cell sorting (FACS), DNA replication by EdU incorporation, G1 versus G2 distribution by aurora B staining, and for chromatin morphology and cohesin localization by DAPI and SCC1 staining, respectively (Fig 5A). The Hi-C biological replicates were highly consistent (Appendix Table S1 and Fig S5), and the low frequency of inter-chromosomal interactions, only representing 1.5% for prometaphase and between 4-6% for all other Hi-C libraries, was indicative of high-quality data sets (Appendix Table S1).

**Figure 5.**
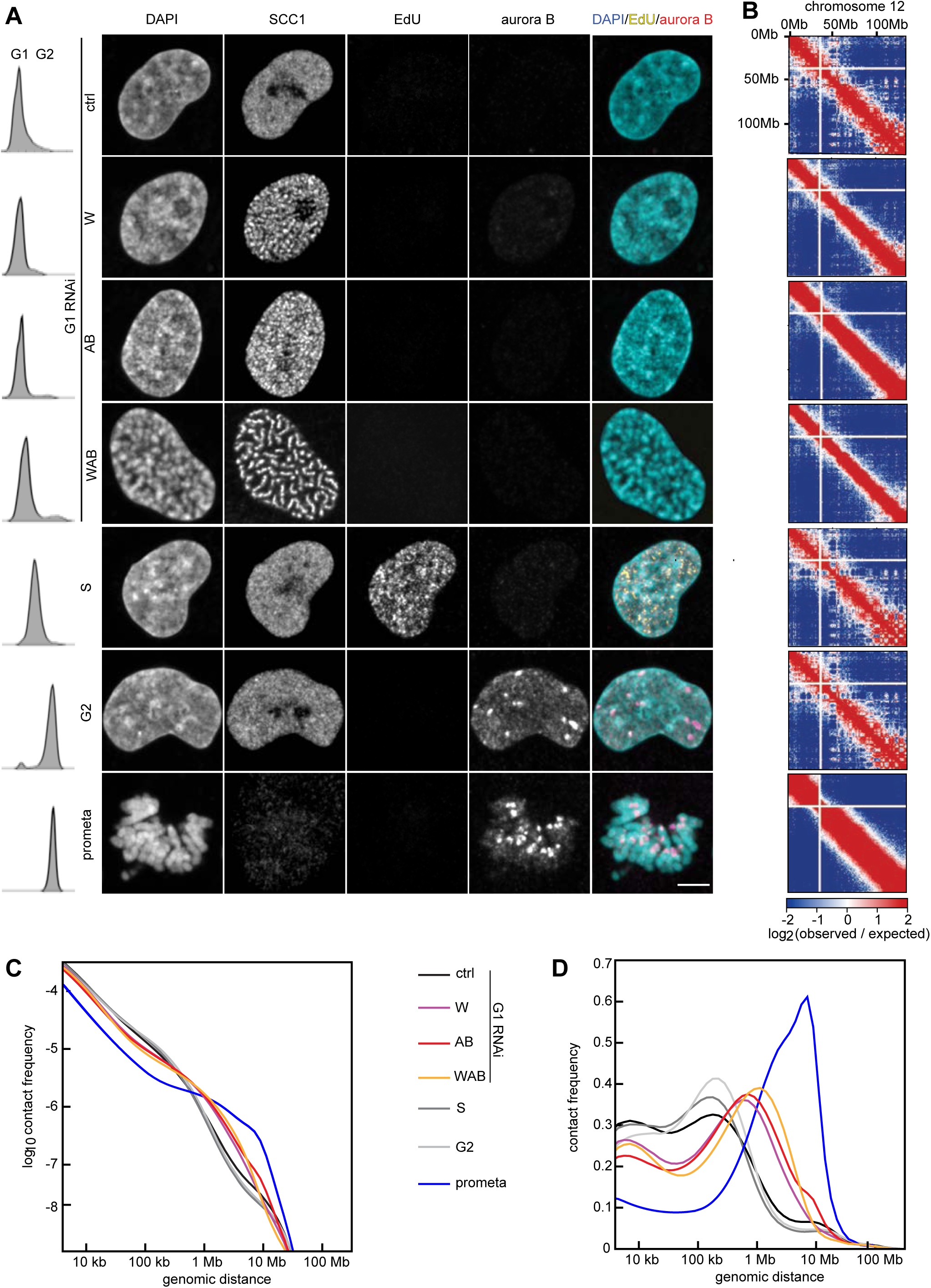
Hi-C of RNAi WAPL and PDS5A/B depletion shows longer-range contacts. A. Immunofluorescence staining of DAPI, SCC1, EdU, Aurora B and joint DAPI/EdU/Aurora B staining (from left to right) for control G1-phase (ctrl), WAPL depleted (W), PDS5A/PDS5B depleted (AB), joint WAPL/PDS5A/PDS5B depleted (WAB), S-phase (S), G2-phase (G2) and pro-metaphase (Prometa) cells (from top to bottom) together with flow-cytometry profiles on the left. Scale bar shows 5 μm. B. Coverage-corrected Hi-C contact enrichment matrices (using HOMER) of chromosome 12, for the same conditions as in (A). C,D. Intra-chromosomal contact frequency distribution as a function of genomic distance using equal sized (C) or logarithmically increasing (D) genomic distance bins, for control G1-phase (black), WAPL depleted (purple), PDS5A/PDS5B depleted (red), joint WAPL/PDS5A/PDS5B depleted (WAB), S-phase (dark grey), G2-phase (light grey) and pro-metaphase (blue) cells.

Coverage-corrected Hi-C contact matrices in cells depleted of WAPL, PDS5A and PDS5B, and all three proteins showed increasing similarities to matrices from prometaphase cells (Fig 5B and Fig S5A and see below). Compared to control G1 cells, contact frequencies were reduced in the 50-250 kb range but increased in the 0.4-2.5 Mb range, with a peak around 600-700 kb in cells depleted of WAPL or PDS5A and PDS5B, and around 1.1 Mb in cells depleted of all three proteins (Figs 5C and D; Appendix Figs S5B and C).

Compartmentalization was reduced in cells depleted of WAPL, PDS5A and PDS5B, or all three proteins, with WAPL depleted cells showing the weakest and triply depleted cells showing the strongest reduction (Fig 6A and Appendix Table 1). The compartment signal derived by *Eigenvector* analysis from Hi-C data confirmed this, as the pattern of alternating A and B compartments observed in control cells was partially lost following depletion of either WAPL or PDSD5A and PDS5B, and was even more strongly reduced in triply depleted cells (Fig 6A, top panels above the Hi-C maps). The compartment signals weakened genome-wide, as could be seen by linearly fitting the signals detected in experimental cells to those obtained in G1 control cells. This resulted in slope values of 0.5, 0.34 and 0.26 for cells depleted of WAPL, PDS5A and PDS5B, or all three proteins, respectively (Fig 6B), confirming that WAPL depletion reduces compartmentalization the least and depletion of all three proteins the most. A strong reduction in compartmentalization was also seen in genome-wide aggregate contact enrichment analyses of 50 compartment categories (Fig 6C).

**Figure 6.**
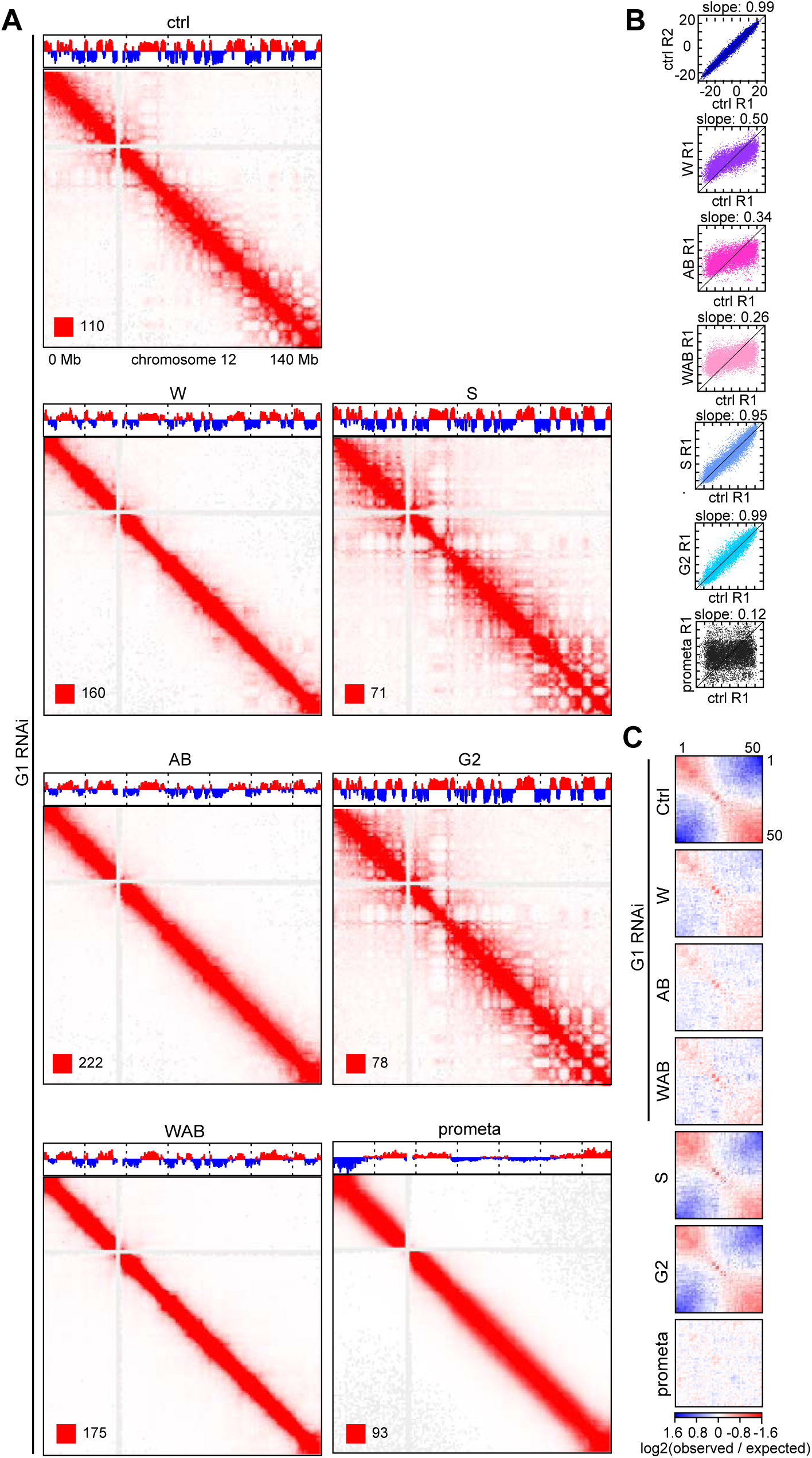
Compartmentalisation weakens upon RNAi WAPL and PDS5A/B depletion. A. Coverage-corrected Hi-C contact matrices of chromosome 12, for control-depleted (ctrl), WAPL depleted (W), PDS5A/PDS5B G1-phase (AB), joint WAPL/PDS5A/PDS5B depleted (WAB) cells in G1 phase, as well as S-phase (S), G2-phase (G2) and pro-metaphase (Prometa) cells. The corresponding compartment signal tracks at 250kb bin resolution are shown above the matrices. The matrices were plotted using Juicebox. B. Scatter plots of compartment signal values at 250kb bin resolution, in the second replicate of control-depleted G1-phase cells (ctrl R2) and the first replicates (R1) of the other conditions (vertical axes) compared to the first replicate of control-depleted G1-phase cells (ctrl R1, horizontal axis). The slope of the linear fit is shown above the plots, and x=y (slope=1) is drawn as black lines. C. Inter-chromosomal contact enrichment between genomic bins with varying compartment signal strength from most B-like (1) to most A-like (50), for the same conditions as in (A).

Whereas compartmentalization was reduced in cells depleted of WAPL, PDS5A and PDS5B, strikingly, many TADs were larger in size than in control cells (Fig 7A; interactions detected in the indicated experimental conditions are shown in the upper right halves of the spilt Hi-C maps; interactions that are specifically occurring in the experimental conditions but not in G1 control cells are shown in difference maps in the lower left halves). Whereas in control cells only relatively few interactions outside of TADs could be observed, these became more prominent in cells depleted of WAPL, PDS5A and PDS5B, resulting in the partial fusion of typically two or three TADs into one. This phenomenon resulted in a decrease in TAD numbers (Fig 7B), an increase in TAD size (Fig 7C), and a reduction in the insulation between TADs (Fig 7D) at relatively constant genome coverage by TADs (Appendix Table 1). These effects were weakest in cells depleted only of WAPL and most pronounced in cells depleted of all three proteins.

**Figure 7.**
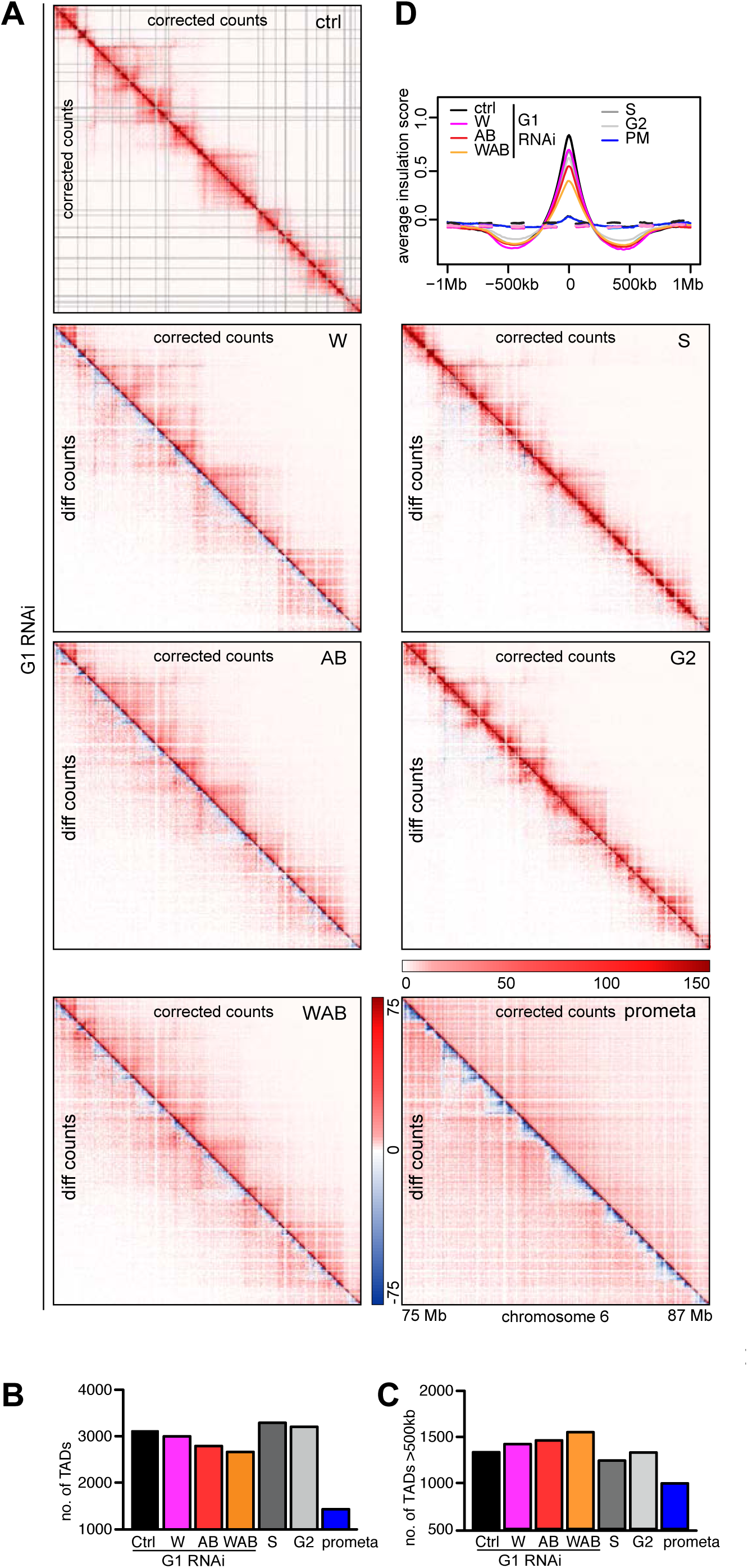
TAD boundaries weaken upon RNAi WAPL and PDS5A/B depletion. A. Coverage-corrected Hi-C contact count matrices (upper triangles, using HOMER) in the 75-87Mb region of chromosome 6, for control-depleted (ctrl), WAPL depleted (W), PDS5A/PDS5B depleted (AB), joint WAPL/PDS5A/PDS5B depleted (WAB) cells in G1 phase, as well as S-phase (S), G2-phase (G2) and pro-metaphase (Prometa) cells. Contact count difference matrices compared to ctrl-G1 (lower triangles) are shown for all non-control conditions. B-C. The number of TADs (B) and the number of long (>500kb) TADs (C) in the same conditions as in (A). Colours are as in Fig 5C. D. Average insulation score around TAD boundaries identified in G1 control HeLa cells, for the same conditions as in (A).

These results indicate that depletion of WAPL, PDS5A and PDS5B suppresses A and B compartments and enables the formation of long-range interactions that extend beyond the borders of TADs as they occur in control cells, resulting in the partial fusion of TADs.

### Chromatin looping is altered differently following depletion of WAPL and PDS5 proteins

Interestingly, close inspection of the interaction matrices revealed that point interactions such as those at the corners of TADs were altered differently upon depletion of WAPL and PDS5 proteins (Fig 8; for examples of point interactions called by “juicer-tool” see lower left half of the split Hi-C matrices in Fig 8A, marked by small black dots).

**Figure 8.**
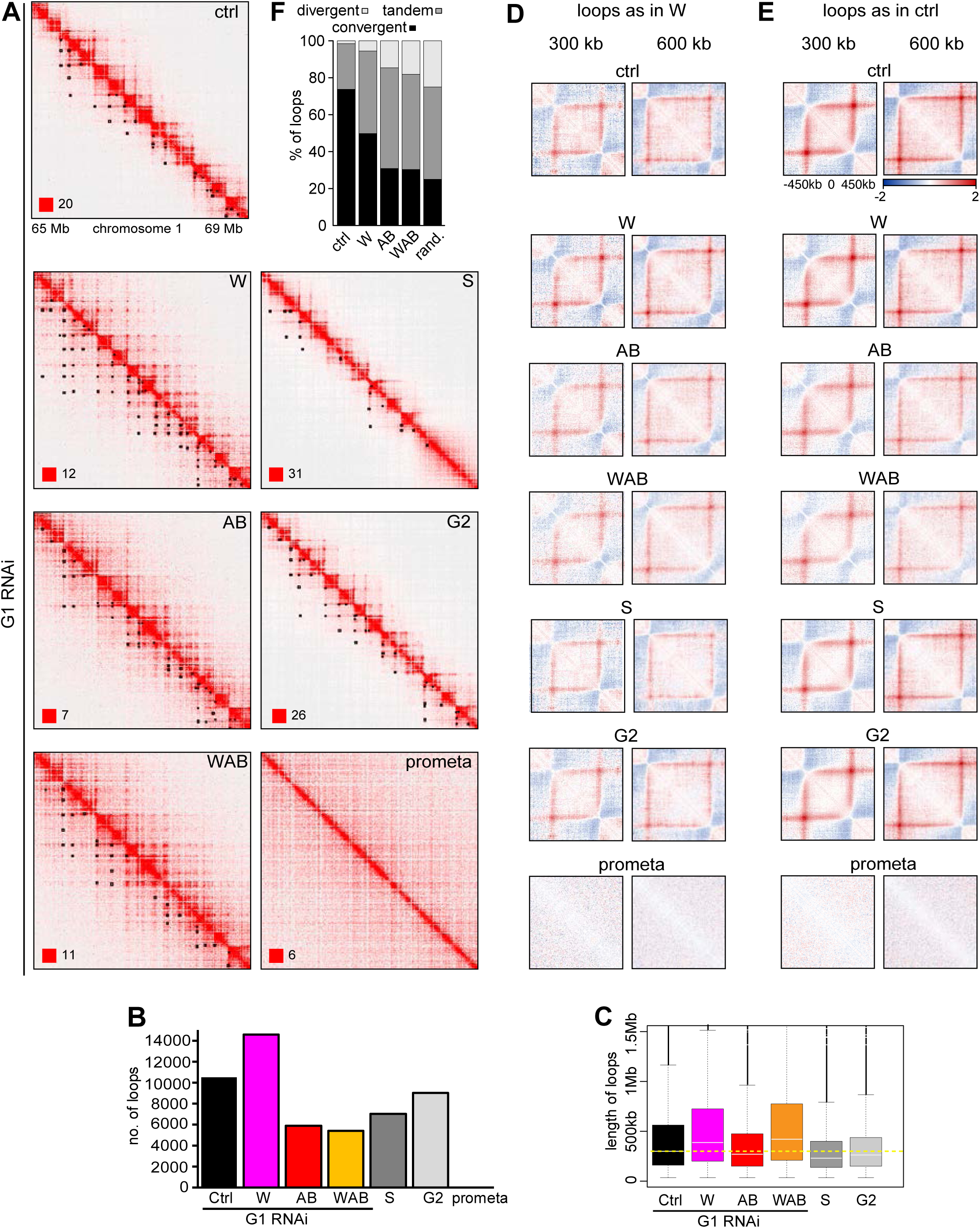
RNAi WAPL and PDS5A/B depletion affects chromatin loops differently. A. Coverage-corrected Hi-C contact matrices of the 65-69 Mb region of chromosome 1, for control-depleted (ctrl), WAPL depleted (W), PDS5A/PDS5B depleted (AB) and joint WAPL/PDS5A/PDS5B depleted (WAB) cells in G1 phase, as well as S-phase (S), G2-phase (G2) and pro-metaphase (Prometa) cells. Loops identified in this region are marked by black circles in the upper triangle. The matrices were plotted using Juicebox. B. Number of loops in the same conditions as in (A). Colours are as in Fig 5C. C. Loop length distribution in the same conditions as in (A). Colours are as in (B). D. Average contact enrichment around the 207 x 300 kb long and 82 x 600 kb long loops identified in WAPL depleted cells but not in G1 control HeLa cells, for the same conditions. E. Average contact enrichment around the 207 x 300 kb long and 82 x 600 kb long loops identified in G1 control HeLa cells, for the same conditions. F. The proportion of convergent (black), tandem (dark grey) and divergent (light grey) CTCF binding orientation for loops with both anchors overlapping SMC3 and CTCF ChIP-seq peaks as well as unambiguous CTCF binding directions, for loops identified in control-depleted cells in G1 phase (ctrl, 2,691 loops), in WAPL depleted but not in control-depleted cells in G1 phase (WAPL, 2,255 loops), in PDS5A/B depleted but not in control-depleted cells in G1 phase (AB, 628 loops) and joint WAPL/PDS5A/PDS5B depleted but not in control-depleted cells in G1 phase (WAB, 769 loops). The theoretically expected random proportions assuming no directionality bias are shown as comparison (rand.).

In WAPL depleted cells, both the number of these loops and their length were increased from 10,310 in control cells (median length 262.5 kb) to 14,557 (median length 387.5 kb; Figs 8B and C; note that in this experiment more loops could be identified than in Appendix Fig S2C because more replicates were analyzed). Following WAPL depletion, more of the looping loci (loop anchors) participated in multiple interactions (55.4%) than in control cells (40.7%; Appendix Fig S6A left panel), resulting in “chains” and “networks” of loops (examples of loop networks can be seen in Fig 8A in the second panel from the top as parallel chains of dots). In the case of networks, 21.6% of these contained at least 10 loops in WAPL depleted cells, compared to only 6.4% in control cells (Appendix Fig S6A right panel). However, aggregate loop analysis revealed that, on average, the loops specifically detected in WAPL depleted cells also existed in control cells, albeit below the detection threshold (Fig 8D, top two panels). In contrast, WAPL depletion did not strengthen the contact enrichment of the looping loci that were already above detection threshold in control cells (Fig 8E, top four panels). These observations indicate that the loops that can be detected specifically in WAPL depleted cells are formed by extension of shorter loops that are also present in control cells, and that the extended loops in WAPL depleted cells also exist in control cell populations, but at much lower frequencies than after WAPL depletion. In other words, the same set of interactions exist in control and WAPL depleted cells, but long-range interactions are much more frequent after WAPL depletion.

Unexpectedly, loops were affected differently in cells in which both PDS5 proteins were depleted. These contained many fewer loops than control cells, both when depleted alone (5,893) or when co-depleted with WAPL (5,259; Fig 8B), and the contact enrichment of looping loci that were present in control cells were weakened (Fig 8E panels in third and fourth row). This is surprising given that genome-wide, contact frequencies were altered very similarly in cells depleted of WAPL and of PDS5 proteins (Figs 5C and D; Appendix Figs S5B and C). The situation in cells depleted of PDS5 proteins is, therefore, reminiscent of the situation in CTCF depleted cells, in which loop numbers are also greatly reduced, whereas interaction frequencies at the genomic level remain unchanged (Figs 2F and I). This similarity raises the interesting possibility that PDS5 proteins are required for the boundary function of CTCF (see Discussion).

### Loops specifically detected in cells depleted of PDS5 proteins and WAPL “violate” the CTCF convergence rule

Remarkably, depletion of PDS5 proteins also led to a strong violation of the CTCF convergence rule (Fig 8F). To be able to determine this, we performed chromatin immunoprecipitation-sequencing experiments (ChIP-seq) to identify the genomic distribution of cohesin and CTCF in control cells, and in cells depleted of WAPL, or PDS5A and PDS5B, or all three proteins (Appendix Fig S6B). To analyze the orientation of CTCF sites at loop anchors, we identified contacts that overlapped with both cohesin (SMC3) and CTCF ChIP-seq peaks, and had a consensus CTCF binding motif. In control cells, 73.8% of loops fulfilling these criteria were anchored at convergent CTCF sites, whereas many fewer had been formed between tandemly oriented (24.7%) and divergent (1.5%) sites (Fig 8F), confirming previous reports (Rao *et al*, 2014; Vietri Rudan *et al*, 2015). However, of the loops that fulfilled these criteria and were specifically detected in cells depleted of PDS5A and PDS5B (i,.e., not detectable in control cells), only 30.9% of loops were anchored by convergent CTCF sites. Instead, most of them (54.5%) were anchored by tandemly oriented CTCF sites, and also many more than normally by divergent sites (14.6%). This distribution is much more similar to what would be expected if CTCF sites were oriented randomly at loop anchors (25% convergent, 50% tandem, 25% divergent) than to the distribution seen in control cells.

An even more randomized distribution of CTCF site orientations was observed at the anchors of loops specifically detected in cells co-depleted of both PDS5 proteins and WAPL (30.4% convergent, 51.5% tandem and 18.1% divergent), whereas loops specifically detected in cells depleted of WAPL alone violated the convergence rule to a lesser extent (49.8% convergent, 44.7% tandem and 5.5% divergent, Fig 8F). Interestingly, in the “chains” of loops, in which the upstream anchor of a loop is also the downstream anchor of a consecutive loop of the chain, the CTCF convergence rule was more frequently violated by internal loops than by the loops which were formed by the outmost anchors (Appendix Fig S6D). This contrast was particularly strong in WAPL depleted cells, while remarkably, the favored CTCF orientation was reversed upon joint depletion of both PDS5 proteins and WAPL.

These results indicate that in cells depleted of PDS5 proteins and WAPL, CTCF boundaries can be skipped more frequently than normally, and that loops which are not anchored by convergent CTCF sites can be formed much more frequently.

### The effects of WAPL, PDS5A and PDS5B depletion on chromatin structure depend on cohesin and are rapidly reversible

To test if chromosome condensation and altered genomic interactions in cells depleted of WAPL, PDS5A and PDS5B depend on cohesin, we generated cells in which chromatin could be imaged while cohesin was inactivated by auxin mediated degradation. For this purpose we stably expressed the histone H2B fused to mRaspberry in SCC1-GFP-AID cells, co-depleted WAPL, PDS5A and PDS5B by RNAi, and subsequently induced SCC1 degradation (Fig 9A). Also under these conditions, in which most cohesin is bound to chromatin, SCC1-GFP-AID was degraded rapidly in 20 min, leading to the disappearance of vermicelli (Fig 9A right panel). Interestingly, chromosomes de-condensed concomitant with cohesin degradation. Quantification of this phenomenon by automated image segmentation revealed that chromosomes began to de-condense rapidly during the initial 10-20 min, i.e. when cohesin degradation occurred, but continued to expand at a slower rate for at least 60 min (Fig 9B right panel).

**Figure 9.**
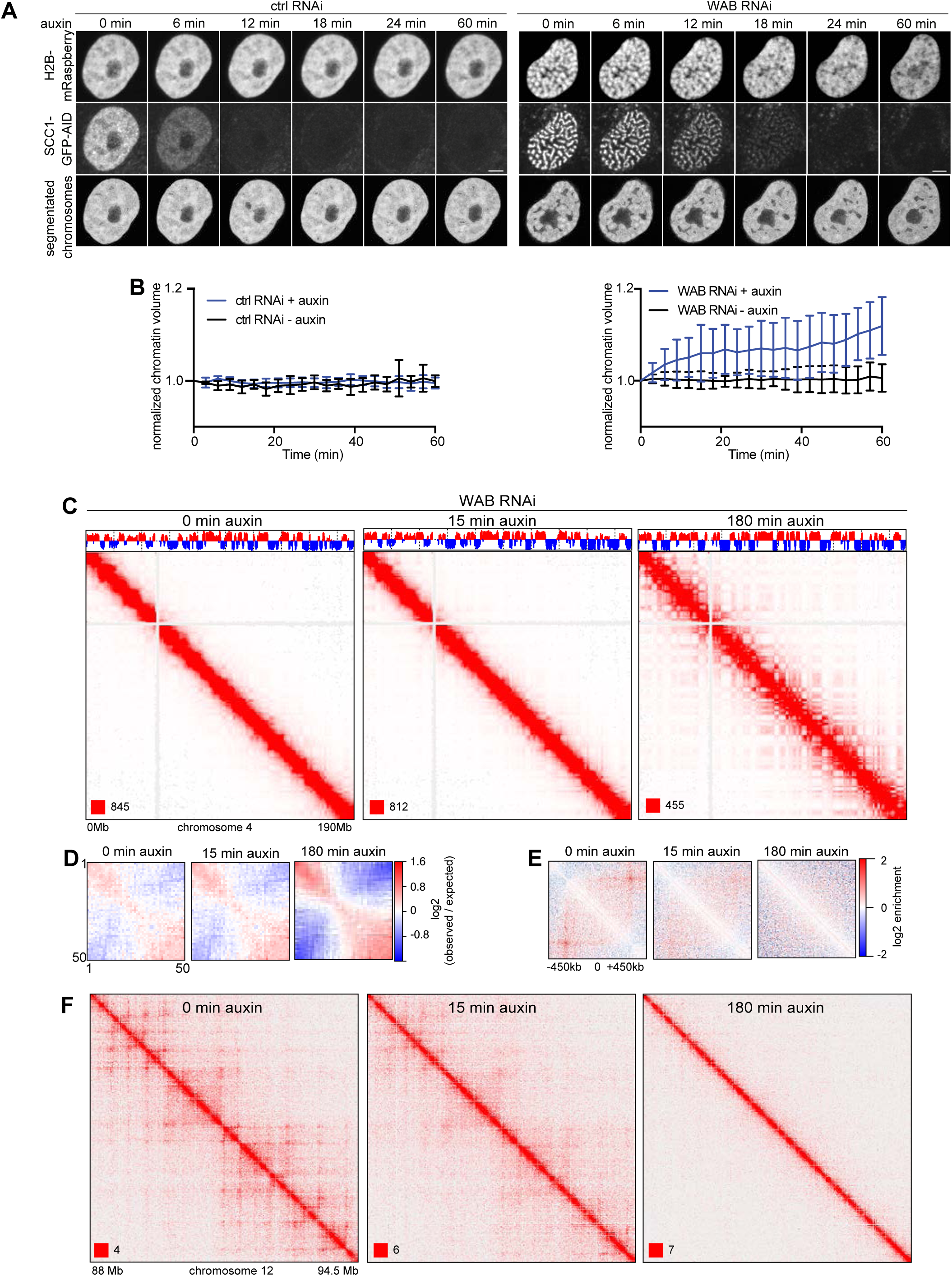
SCC1 degradation in RNAi WAPL and PDS5A/B depleted cells. A. Analysis of the chromosome morphological changes after auxin addition in control-depleted and WAPL/PDS5A/B depleted SCC1-GFP-AID cell lines that stably express mRaspberry-H2B by live cell imaging. Scale bar indicates 5 μm. B. Chromatin volumetric changes in auxin treated and un-treated cells that had been depleted for control (left) and WAB (right). Chromatin volumes are normalized to the chromatin volumes at time 0 and plotted over time (n > 7 in each condition). Error bar depicts standard deviations of the means. C. Coverage-corrected Hi-C contact matrices of chromosome 4, at 0 (left), 15 (centre) and 180 minutes (right) after auxin addition to the WAPL/PDS5A/PDS5B depleted SCC1-AID cell line. The matrices were plotted using Juicebox. The corresponding compartment signal tracks at 250kb bin resolution are shown above the matrices. D. Inter-chromosomal contact enrichment between bins with varying compartment signal strength from most B-like (1) to most A-like (50), for the same conditions as in (C). E. Average contact enrichment around loops, for the 82 x 600 kb long loops identified in G1 control cells. The matrices are centered (0) around the halfway point of the loop anchor coordinates. F. Coverage-corrected Hi-C contact matrices in the 88-94.5 Mb region of chromosome 12, for the same conditions as in (C). The matrices were plotted using Juicebox.

To analyze chromatin structure, we performed Hi-C experiments after auxin treatment for 0, 15 and 180 min. Also in these cells, depletion of WAPL, PDS5A and PDS5B strongly suppressed compartmentalization but increased the size of TADs dramatically. However, addition of auxin reverted these phenotypes, leading after 180 min to interaction matrices that were similar to those seen in cells partially depleted of cohesin, i.e compartments were strengthened (Figs 9C and D), whereas loops and TADs disappeared (Figs 9E and F, Appendix Figs S6E and F).

These results indicate that the strong chromosome condensation seen in cells co-depleted of WAPL, PDS5A and PDS5B depends on cohesin, as does the mild chromatin compaction caused by WAPL depletion in MEFs (Tedeschi *et al*, 2013). Furthermore, these experiments show that also the suppression of compartmentalization and the weakening of TAD insulation in cells depleted of WAPL, PDS5A and PDS5B depend on cohesin, and interestingly they raise the possibility that cohesin inactivation affects different steps of chromosome de-condensation with different kinetics.

### Chromatin structure changes in WAPL, PDS5A and PDS5B depleted cells are similar but not identical to changes that occur during mitotic chromosome condensation

Because co-depletion of WAPL, PDS5A and PDS5B resulted in chromatin condensation reminiscent of that observed in prophase (Fig 3) we compared the Hi-C interaction matrices of these cells with those of cells synchronized in different cell cycle stages, including prometaphase (Fig 5).

A comparison of the different stages showed overall similar genome-wide interaction frequencies in G1, S and G2, with a small increase in short-range interactions (60-800 kb) and a corresponding decrease in long-range interactions (3-30 Mb) in cells in S and G2 relative to G1 (Figs 5C and D). Also compartmentalization (Figs 6A-C) and position, number, size and insulation of TADs (Figs 7A-D) were similar, whereas the number and length of loops decreased in S and increased again in G2, although not to the extent observed in G1 (Figs 8A-C). It will, therefore, be interesting to test if DNA replication interferes with formation or maintenance of loops. As in G1, most loops formed between convergent CTCF sites in S and G2 (75.7% and 71.7% respectively).

In contrast, in prometaphase cells most interactions <0.8 Mb were lost and long-range interactions (0.8-70 Mb) greatly increased, with 8-9 Mb interactions having the highest frequency (Figs 5C and D). Compartments could not be detected (Fig 6A-C), TADs were greatly reduced in number and size (Figs 7B and C), and almost all loops disappeared (Fig 8A-E; “juicer-tool” could only identify 57). These results confirm and extend previous analyses of cell populations in G1, S and prometaphase (Naumova *et al*, 2013) and single-cell analyses of all four cell cycle stages (Nagano *et al*, 2017).

Interestingly, at two levels of genome organization cells depleted of WAPL, PDS5A and PDS5B were in an intermediate state between interphase and mitotic cells. Whereas in interphase and prometaphase cells interactions were most frequently bridging distances of 0.3 and 8-9 Mb, respectively, in cells depleted of WAPL, PDS5A and PDS5B the most frequent interactions occurred between loci 1-1.1 Mb apart (Figs 5C and D). Similarly, compartment “strength” was strongest in interphase cells (slope values of 0.95-0.99), weakest in prometaphase (slope value 0.12), and intermediate in cells depleted of WAPL, PDS5A and PDS5B (slope values 0.26-0.5; Fig 6B). A similar trend was observed for the number of TADs, which decreased following depletion of WAPL, PDS5A and PDS5B, i.e. became more similar also in this respect to prometaphase cells (Fig 7B). These results indicate that depletion of WAPL, PDS5A and PDS5B induces chromosome condensation by changes in genome organization that are similar to those occurring in mitosis, even though the former are caused by cohesin and the latter by condensin.

## Discussion

Although cohesin was initially discovered as a protein complex essential for sister chromatid cohesion, it has long been suspected that cohesin has distinct roles in gene regulation and that it may be defects in these functions and not in cohesion that can contribute to tumorigenesis and lead to developmental defects, called cohesinopathies (reviewed in (Watrin *et al*, 2016)). It has also been realized that cohesin’s function in gene regulation is related to CTCF, as this DNA binding protein recruits cohesin to specific sites in mammalian genomes and depends on cohesin for its ability to mediate gene regulation (Parelho *et al*, 2008; Wendt *et al*, 2008; Busslinger *et al*, 2017). Because CTCF affects gene expression by chromatin looping (Kurukuti *et al*, 2006; Splinter *et al*, 2006), it has been speculated that CTCF and cohesin have discrete functions in chromatin organization, with CTCF specifying the genomic sites at which cohesin functions, and cohesin entrapping DNA sequences in *cis* to generate chromatin loops (Wendt *et al*, 2008; Wendt & Peters, 2009). This model predicts that chromatin loops depend on cohesin but not necessarily on CTCF, whereas conversely the identity of DNA sequences that form loop anchors would depend on CTCF but not on cohesin.

Earlier support for a role of cohesin in chromatin organization came from the observation that cohesin can compact chromatin when cohesin’s residence time and levels are artificially increased on chromatin by depletion of WAPL (Tedeschi *et al*, 2013). However, whether cohesin is actually required for chromatin organization under physiological conditions has remained less clear as earlier cohesin inactivation experiments either only analyzed individual chromatin interactions (Hadjur *et al*, 2009; Mishiro *et al*, 2009; Nativio *et al*, 2009) or did not reveal major changes in TADs and none in compartments ((Seitan *et al*, 2013; Sofueva *et al*, 2013; Zuin *et al*, 2014); chromatin loops were not yet analyzed genome-wide in these studies). Our results reported here remove this uncertainty as they show clearly that TADs and loops cannot be detected in the absence of cohesin. Our experiments further indicate that CTCF is not essential for long-range interactions *per se*, although we can formally not exclude that small amounts of CTCF persisted in our experiments and contributed to the chromatin interactions that were still detectable in CTCF depleted cells. Our results show clearly, however, that CTCF is required for TAD insulation and the detectability of loops.

While cohesin had been predicted to mediate formation of TADs and loops, it was unexpected that cohesin depletion would increase the detectability of A and B compartments, as these are thought to represent different compaction states of chromatin (euchromatin and heterochromatin, respectively) that are generated by post-translational histone modifications which modulate short-range interactions between nucleosomes (Lieberman-Aiden *et al*, 2009; Rao *et al*, 2014). The notion that cohesin is a key determinant of chromatin structure, not only at the level of TADs and loops, but also at the level of compartments, is also supported by our observation that stabilization of cohesin on chromatin by WAPL depletion does not only increase the frequency with which cells can form long chromatin loops but also decreases the detectability of compartments. It will, therefore, be interesting to analyze whether cohesin-mediated long-range interactions affect local nucleosomal interactions, or whether cohesin has a more direct role specifying euchromatic and heterochromatic regions.

We suspect that our results are at variance with previous cohesin and CTCF inactivation experiments in which only mild phenotypes could be detected (Seitan *et al*, 2013; Sofueva *et al*, 2013; Zuin *et al*, 2014) because these proteins may have been depleted to various degrees in the different studies. This possibility is consistent with our observation that TAD patterns seem unaffected in SCC1-GFP-AID and CTCF-GFP-AID expressing cells if these are not treated with auxin (Appendix Fig S2D), even though the analysis of chromatin interactions genome-wide (Appendix Fig S2A) clearly indicates that these cells are hypomorphic for the roles of cohesin and CTCF in chromatin organization. It may be difficult to observe major changes in Hi-C experiments in which cohesin and CTCF were depleted incompletely, either because no calibration methods are presently available for Hi-C assays, and/or because small amounts of these proteins are sufficient for their function, as is known to be the case for the cohesion function of cohesin (Heidinger-Pauli *et al*, 2010).

Our results are, however, consistent with several other studies which were deposited on *bioRxiv* (Schwarzer *et al*, 2016; Rao *et al*, 2017) or published in peer-reviewed journals (Nora *et al*, 2017; Haarhuis *et al*, 2017) during preparation of our manuscript. In these studies, chromatin structure was analyzed by Hi-C experiments following inactivation of either cohesin (Rao *et al*, 2017), the cohesin loading complex (Schwarzer *et al*, 2016), CTCF (Nora *et al*, 2017) or WAPL (Haarhuis *et al*, 2017). The results of these and our experiments, combined with the previous observations that cohesin and CTCF are enriched at TAD boundaries and loop anchors (Dixon *et al*, 2012; Rao *et al*, 2014), strongly support the hypothesis that cohesin is a key determinant of chromatin structure at multiple levels, that in the case of TADs and loops cohesin has a direct role in mediating long-range interactions, possibly by entrapping DNA sequences in *cis*, and that CTCF specifies where in the genome cohesin performs these functions.

Importantly, our results are also fully consistent with the loop extrusion hypothesis (Nasmyth, 2001; Sanborn *et al*, 2015; Fudenberg *et al*, 2016) based on which it had been predicted that WAPL depletion would lead to the formation of extended chromatin loops (Fudenberg *et al*, 2016). The fact that more and larger loops can indeed be detected in WAPL depleted cells (Figs 8B and C and (Haarhuis *et al*, 2017)) and that also these depend on cohesin (Figs 8B and C) confirms this prediction and indicates that WAPL depletion causes chromatin compaction and accumulation of cohesin in axial positions (vermicelli) by loop extension. As had been pointed out by (Fudenberg *et al*, 2016), the generation of these loops could be enabled by the increase in cohesin’s residence time that WAPL depletion is known to cause (Tedeschi *et al*, 2013), which would allow cohesin to move along chromatin fibers with higher processivity. In addition, the increase in cohesin’s copy number on chromatin that results from its increased residence time could also contribute to the frequency with which extended loops can be detected in WAPL depleted cells, because it is possible that in control cells at steady state not all pairs of *loci* that could form loops are actually occupied by cohesin.

Interestingly, our analyses indicate that the shorter loops detected in control cells persist after WAPL depletion (Fig 8D). Conversely, also the extended loops seen in WAPL depleted cells can on average be detected in aggregate loop analyses even in control cells (Fig 8E), even though these extended loops cannot be identified in these cells individually because they are below the detection limit. These observations imply that WAPL depletion does not lead to the formation of loops that cannot exist in control cells, but rather increases the frequency with which extended loops exist in the cell population.

WAPL is known to bind PDS5A and PDS5B (Kueng *et al*, 2006). Like WAPL, these proteins have been implicated in releasing cohesin from DNA (Rowland *et al*, 2009; Shintomi & Hirano, 2009; Ouyang *et al*, 2016), but PDS5 proteins also have positive functions in cohesion (for references see (Hons *et al*, 2016)) and *in vitro* can promote loading of cohesin onto DNA (Murayama & Uhlmann, 2015). In agreement with a recent report (Ouyang *et al*, 2016), we found that depletion of PDS5 proteins can also increase cohesin’s residence time on chromatin, although not to the same extent as depletion of WAPL, and cause chromatin compaction and cohesin accumulation in vermicelli. Despite these phenotypic similarities, which are consistent with the idea that WAPL can release cohesin from DNA more efficiently in the presence of PDS5 proteins than in their absence, depletion of PDS5 proteins caused local changes in Hi-C interactions patterns that were strikingly different from those seen after WAPL depletion. Whereas WAPL depletion increased the number of detectable loops by 41.2 %, co-depletion of PDS5A and PDS5B reduced loop numbers by 42.8 % (Fig 8B). This effect of PDS5 inactivation was dominant over the effect of WAPL depletion, as depletion of all three proteins reduced loop numbers by 49.0 %. Despite these differences, depletion of WAPL and PDS5 proteins increased long-range interactions genome-wide in very similar ways (Figs 5C and D), resulting in the detectability of slightly fewer but larger TADs (Figs 7B and C) and, as mentioned, chromatin compaction and vermicelli formation. These results imply that PDS5 proteins are not required for long-range chromatin interactions *per se*, but for their precise location. In other words, cells depleted of WAPL and PDS5 proteins may have similar number of loops, including extended loops, but in the former case they might be formed more frequently between well-defined pairs of sequences, whereas in the latter there may be more heterogeneity between loci that function as loop anchors, which would prevent their detection as specific corner peaks (i.e. loops) in Hi-C matrices. This particular phenotype caused by depletion of PDS5 proteins is similar to the effect of CTCF depletion, which also reduces the number of detectable loops (Fig 2I) without affecting the total number of long-range chromatin interactions genome wide (Fig 2F). This phenotypic similarity raises the interesting possibility that the ability of CTCF to function as a boundary element in loop extrusion depends on PDS5 proteins.

How PDS5 proteins could perform this function is unclear, as is the mechanistic basis of CTCF’s boundary function, but one possibility emerges from the previous observations that the cohesin loading complex stimulates cohesin’s ATPase activity (Murayama & Uhlmann, 2014), is required for vermicelli formation (Haarhuis *et al*, 2017) and competes with PDS5 proteins for cohesin binding (Kikuchi *et al*, 2016). These findings would be consistent with the possibility that loop extrusion is mediated by cohesin’s ATPase activity following its stimulation by the loading complex (as discussed in a recent *bioRxiv* pre-print by (Rhodes *et al*, 2017)), and that PDS5 proteins would inhibit this activity. If correct, our data imply that PDS5 proteins might mediate this effect in a CTCF dependent manner. A different, but equally important question will be to understand how CTCF containing boundaries can be skipped to form extended loops, a phenomenon that according to our data must occur occasionally in control cells and more frequently in cells depleted of WAPL and PDS5 proteins and that leads to violation of the CTCF convergence rule and, in cells depleted of WAPL and PDS5 proteins, to chromatin compaction and formation of vermicelli.

## Materials and Methods

### Generation of cell lines for auxin mediated degradation

HeLa Kyoto cells were cultured as described previously (Nishiyama *et al*, 2010). The Hela Kyoto cell line C-terminally-tagged with a SCC1 auxin-inducible degron (AID) was created by CRISPR/Cas9 mediated genome editing using a double-nickase strategy (Ran *et al*, 2013). The homologous recombination template used introduced the sequences coding for monomeric EGFP (L221K) and the *Arabidopsis thaliana* IAA1771-114 (AID*) mini-degron (Morawska & Ulrich, 2013). Single clones were selected by flow cytometry and confirmed to be homozygous by PCR. One of these clones was transduced with lentiviruses containing Tir1 from rice (*Oryza sativa*) using the plasmid pRRL-SOP (for SFFV, OsTir1, Puro), which comprises the constitutive promoter from spleen focus forming virus followed by OsTir1-3x-Myc-tag-T2A-Puro. The gRNAs sequences that were used for generating the SCC1 auxin-inducible degron cell line were gRNA1: CACCGCAAATTGCCCCCATGTGTA, and gRNA2: CACCGCTTATATAATATGGAACCT. Primers used for genotyping were: forward primer: TCAGGTATTGCCGCTGTTGT, and reverse primer: GTGTGCAGCACTGGAAAAGG. The CTCF-GFP-AID cell line was generated in the same way with gRNA sequences gRNA1: TCAGCATGATGGACCGGTGATGG, and gRNA2: TGAGGATCATCTCGGGCGTGAGG. The primers used for genotyping were: forward primer: ACTGTTAATGTGGGCGGGTT and reverse primer: CTGGGTGCCATTCTGCTACA.

### Synchronization protocol

Cells were synchronized at the G1/S phase boundary by two consecutive arrest phases with 2 mM thymidine and released into fresh medium for 2h for (S phase), 6 h (G2 phase) or 15 h (G1 phase). For prometaphase arrest, cells were released from double thymidine block and 8 h later Nocodazole was added at the concentration of 100 ng/ml. Five hours later, PM cells were removed by shake off. For each cell cycle stage, cells were fixed for Hi-C, microscopy and FACS, respectively. Cell cycle profiling was performed using propidium iodide staining (Ladurner *et al*, 2014).

### RNA interference

For RNAi experiments, the cells were transfected with siRNAs as described previously (van der Lelij *et al*, 2014) just before adding thymidine. The following target sequences of siRNAs (Ambion) were used: WAPL (5′-CGGACUACCCUUAGCACAAtt-3′), PDS5A (5′-GCUCCAUAUACUUCCCAUGtt-3′), PDS5B (5′-GAGACGACUCUGAUCUUGUtt-3′), NIPBL (5’-GCAUCGGUAUCAAGUCCCAtt-3’) and SMC2 (5’-UGCUAUCACUGGCUUAAAUtt-3’). The cells were transfected by incubating duplex siRNA with RNAi-MAX transfection reagent in antibiotic free growth medium at 100 nM. After 72 h RNAi treatment, cells were harvested for the experiments.

### Chromatin fractionation

Cells were extracted in a buffer consisting of 20 mM Tris-HCl (pH 7.5), 100 mM NaCl, 5 mM MgCl, 2 mM NaF, 10% glycerol, 0.2% NP40, 20 mM β-glycerophosphate, 0.5 mM DTT and protease inhibition cocktail (Complete EDTA-free, Roche). Chromatin pellets and supernatant were separated and collected by centrifugation at 2,000 *g* for 5 min. The chromatin pellets were washed three times with the same buffer.

### Antibodies

NIPBL antibodies were a generous gift from Tom Strachan (Seitan *et al*, 2006). MAU2 antibodies were reported previously (Watrin *et al*, 2006). Rabbit anti-CTCF (Peters laboratory A992) was reported previously (Busslinger *et al*, 2017). The following commercial antibodies were used: SMC1 (Bethyl Laboratories, A300-055A), SMC3 (Bethyl Laboratories, A300-060A), SCC1 (EMD Millipore Corporation, 53A303), PDS5A (Bethyl Laboratories, A300-089A), a-tubulin (Sigma-Aldrich, T5168), histone H3 (Santa Cruz Biotechnology, sc-8654), phospho-histone H3 (Ser10) (Cell Signaling, #9701), GFP (Roche, clones 7.1 and 13.1; Abcam, ab13970), Aurora B (BD Transduction Laboratory, 611083). The following secondary antibodies were used: goat anti-rabbit IgG Alexa-Fluor-488 and −568, anti-mouse IgG Alexa-Fluor-488, −568 (all Molecular Probes) for IF and anti-rabbit or mouse Ig, HRP-linked whole antibody (GE Healthcare) and polyclonal rabbit anti-goat or anti-rat immunoglobulins/HRP (Dako) for WB.

### Immunofluorescence

Cells were fixed with 4% formaldehyde for 10 min at room temperature and permeabilized with 0.2% Triton X-100 in PBS for 5 min. Pre-extraction was carried out for 1 min in 0.1% Triton X-100 in PBS before fixation. After blocking in 3% BSA in PBS-T 0.01% for 1 h, the cells were incubated with the primary antibodies overnight at 4 °C and subsequently incubated with the secondary antibodies for 1 hour at room temperature. After counterstaining the DNA by 10 min incubation in 0.1 μM DAPI in PBS, the chambers were perfused with vector shield (Vector Laboratories) before imaging.

### Live cell imaging

SCC1-GFP-AID cells were seeded on Lab-Tek chambered coverslips (Thermo Scientific Nunc) and treated with control or WAPL, PDS5A and PDS5B targeting siRNAs for 72 hours. Before imaging, the culture medium was exchanged for pre-warmed CO_2_-independent medium (Gibco). DNA was visualized by staining with 0.5 μM SiR-DNA (Spirochrome, CHF280.00) or stably expressing mRaspberry-H2B. Imaging was performed with a LSM780 confocal microscope (Carl Zeiss), using a 63×/1.4 numerical aperture (N/A) objective. After treatment with 500 μM auxin, three-dimensional images with 520 × 520 × 35 voxels (voxel size x × y × z: 100 nm × 100 nm × 500 nm) were collected every 3 minutes, or two-dimensional images with 1568 × 1568 pixels (pixel size x × y: 108 nm × 108 nm) every 2 minutes.

### Inverse Fluorescence recovery after photobleaching (iFRAP)

SCC1-GFP cells that stably express DHB-mKate2 and that had been depleted for control, W, AB or WAB were imaged by a LSM780 confocal microscope (Carl Zeiss), using a 63×/1.4 numerical aperture (N/A) objective. Two images were acquired before bleaching half of the nucleus by 100% transmission of a 488 nm laser. The recovery was quantified using the ZEN2011 software, as the difference between the mean fluorescence signal intensity of the bleached and the unbleached regions. G1 phase cells were identified by nuclear localisation of DHB-mKate2 signals (Spencer *et al*, 2013).

### Segmentation of nuclear mass and quantification of volume and intensity distribution

The nuclear mass was segmented in two sequential steps from the mRaspberry-H2B channel using a fully automated script developed in MATLAB (The MathWorks Inc., Natick, MA). In the first step, the nuclear region of interest is detected independently at each time point. In this process, a Gaussian filter of kernel size 3 with standard deviation 1.5 is applied first to each z-slice of original image to reduce the effect of noise. Each of the original z-slices is binarized by combining two threshold values computed adaptively from the slice (2D) and from the entire stack (3D) as described in (Heriche *et al*, 2014). The nuclear mass of interest was detected from the binarized images by connecting component analysis. The nuclear mass detected in the first step contains almost the entire nuclear volume, thus it may contain low intensity regions including nucleoli. In the second step, to detect nucleus structural changes sensitively, the initial nuclear mass is segmented into high and low intensity voxels, of which the high intensity voxels are retained for further analysis. The first time point is segmented in similar fashion using both 2D and 3D threshold values determined adaptively from the histogram constructed from the voxels inside the initial nuclear mass. The total intensity of the re-segmented region in the first frame is taken as a reference to guide the segmentation of the stacks in later time points in order to obtain the same amount of total intensity inside the nuclear mass. Then the volume of the refined segmented nuclear region is used for quantification where the volume in the first time point is normalized to 1. The higher intensity voxels in the initial segmented nuclear mass are also analyzed independently by clustering the voxels into two classes based on their intensity value. In this process, the “first class” consists of the brightest voxels that add up 50% of the total intensity inside the nucleus and the remaining voxels form the “second class”. Next, the volume of each group of voxels is computed and the ratio of the volume of the brighter first class to that of the dimmer second class is used for quantification. The ratio obtained in the first frame is normalized to 1.

### ChIP-seq peak calling and calculation of peak overlaps

Cohesin (SMC3) and CTCF ChIP was performed as described in (Wendt *et al*, 2008). Peaks were called by the MACS algorithm version 1.4.2 (Zhang *et al*, 2008), using a P-value threshold of 1e-10 and by using sample and input read files. We identified sites of overlapping peaks between different conditions as well as between SMC3 and CTCF peaks using the MULTOVL software (Aszódi, 2012). We applied an inclusive type of overlap display (“union”), in which coordinates of overlapping peaks are merged into one common genomic site.

### Hi-C experiments

Hi-C libraries were generated as described in (Nagano *et al*, 2015), with modifications as described below. 3 to 4 x 10_7_ HeLa cells were fixed in 2% formaldehyde (Agar Scientific) for 10 minutes, after which the reaction was quenched with ice-cold glycine (Sigma; 0.125 M final concentration). Cells were collected by centrifugation (400 x g for 10 minutes at 4°C), and washed once with 50 ml PBS pH 7.4 (Gibco). After another centrifugation step (400 x g for 10 minutes at 4°C), the supernatant was completely removed and the cell pellets were immediately frozen in liquid nitrogen and stored at −80°C. After thawing, the cell pellets were incubated in 50 ml ice-cold lysis buffer (10 mM Tris-HCl pH 8, 10 mM NaCl, 0.2% Igepal CA-630, protease inhibitor cocktail (Roche)) for 30 minutes on ice. After centrifugation to pellet the cell nuclei (650 x g for 5 minutes at 4°C), nuclei were washed once with 1.25 x NEBuffer 2 (NEB). The nuclei were then resuspended in 1.25 x NEBuffer 2, SDS (10% stock; Promega) was added (0.3% final concentration) and the nuclei were incubated at 37°C for one hour with agitation (950 rpm). Triton X-100 (Sigma) was added to a final concentration of 1.7% and the nuclei were incubated at 37°C for one hour with agitation (950 rpm). Restriction digest was performed overnight at 37°C with agitation (950 rpm) with *HindIII* (NEB; 1500 units per 7 million cells). Using biotin-14-dATP (Life Technologies), dCTP, dGTP and dTTP (Life Technologies; all at a final concentration of 30 μM), the *HindIII* restriction sites were then filled in with Klenow (NEB) for 75 minutes at 37°C. The ligation was performed for four hours at 16°C (50 units T4 DNA ligase (Life Technologies) per 7 million cells starting material) in a total volume of 5.5 ml ligation buffer (50 mM Tris-HCl, 10 mM MgCl_2_, 1 mM ATP, 10 mM DTT, 100 μg/ml BSA, 0.9% Triton X-100) per 7 million cells starting material. After ligation, crosslinking was reversed by incubation with Proteinase K (Roche; 65 μl of 10mg/ml per 7 million cells starting material) at 65°C overnight. An additional Proteinase K incubation (65 μl of 10mg/ml per 7 million cells starting material) at 65°C for two hours was followed by RNase A (Roche; 15 μl of 10mg/ml per 7 million cells starting material) treatment and two sequential phenol/chloroform (Sigma) extractions. After DNA precipitation (sodium acetate 3 M pH 5.2 (1/10 volume) and ethanol (2.5 x volumes)) overnight at −20°C, the DNA was spun down (centrifugation 3200 x g for 30 minutes at 4°C). The pellets were resuspended in 400 μl TLE (10 mM Tris-HCl pH 8.0; 0.1 mM EDTA), and transferred to 1.5 ml eppendorf tubes. After another phenol/chloroform (Sigma) extraction and DNA precipitation overnight at −20°C, the pellets were washed three times with 70% ethanol, and the DNA concentration was determined using Quant-iT Pico Green (Life Technologies). For quality control, candidate 3C interactions were assayed (primers available upon request) by PCR, and the efficiency of biotin incorporation was assayed by amplifying a 3C ligation product (primers available upon request), followed by digest with *HindIII* or *NheI*.

To remove biotin from non-ligated fragment ends, 30 to 40 μg of Hi-C library DNA were incubated with T4 DNA polymerase (NEB) for 4 hours at 20°C, followed by phenol/chloroform purification and DNA precipitation overnight at −20°C. After a wash with 70% ethanol, sonication was carried out to generate DNA fragments with a size peak around 400 bp (Covaris E220 settings: duty factor: 10%; peak incident power: 140W; cycles per burst: 200; time: 55 seconds). After end repair (T4 DNA polymerase, T4 DNA polynucleotide kinase, Klenow (all NEB) in the presence of dNTPs in ligation buffer (NEB)) for 30 minutes at room temperature, the DNA was purified (Qiagen PCR purification kit). dATP was added with Klenow exo-(NEB) for 30 minutes at 37°C, after which the enzyme was heat-inactivated (20 minutes at 65°C). A double size selection using AMPure XP beads (Beckman Coulter) was performed: first, the ratio of AMPure XP beads solution volume to DNA sample volume was adjusted to 0.6:1. After incubation for 15 minutes at room temperature, the sample was transferred to a magnetic separator (DynaMag-2 magnet; Life Technologies), and the supernatant was transferred to a new eppendorf tube, while the beads were discarded. The ratio of AMPure XP beads solution volume to DNA sample volume was then adjusted to 0.9:1 final. After incubation for 15 minutes at room temperature, the sample was transferred to a magnet (DynaMag-2 magnet; Life Technologies). Following two washes with 70% ethanol, the DNA was eluted in 100 μl of TLE (10 mM Tris-HCl pH 8.0; 0.1 mM EDTA). Biotinylated ligation products were isolated using MyOne Streptavidin C1 Dynabeads (Life Technologies) on a DynaMag-2 magnet (Life Technologies) in binding buffer (5 mM Tris pH8, 0.5 mM EDTA, 1 M NaCl) for 30 minutes at room temperature. After two washes in binding buffer and one wash in ligation buffer (NEB), PE adapters (Illumina) were ligated onto Hi-C ligation products bound to streptavidin beads for 2 hours at room temperature (T4 DNA ligase NEB, in ligation buffer, slowly rotating). After washing twice with wash buffer (5 mM Tris, 0.5 mM EDTA, 1 M NaCl, 0.05% Tween-20) and then once with binding buffer, the DNA-bound beads were resuspended in a final volume of 90 μl NEBuffer 2. Bead-bound Hi-C DNA was amplified with 7 PCR amplification cycles (36 to 40 individual PCR reactions) using PE PCR 1.0 and PE PCR 2.0 primers (Illumina). After PCR amplification, the Hi-C libraries were purified with AMPure XP beads (Beckman Coulter). The concentration of the Hi-C libraries was determined by Bioanalyzer profiles (Agilent Technologies) and qPCR (Kapa Biosystems), and the Hi-C libraries were paired-end sequenced (HiSeq 1000, Illumina) at the Babraham Institute Sequencing Facility and (HiSeqv4, Illumina) at VBCF NGS.

### Hi-C analyses

#### Hi-C data processing and normalization

Raw sequencing reads were processed using the HiCUP pipeline (Wingett *et al*, 2015). Positions of di-tags were mapped against the human genome (hg19), and experimental artifacts, such as circularized reads and re-ligations, were filtered out, and duplicate reads were removed. Library statistics are presented in Appendix Table 1. Aligned Hi-C data was normalised using *HOMER* v4.7 (Heinz *et al*, 2010) and *Juicer tools* v0.7.5 (Durand *et al*, 2016). Using binned Hi-C data, we computed the coverage- and distance-related background in the Hi-C data employing matrix balancing algorithms at 25kb, 250kb, 1Mb (iterative correction by *HOMER*) and 2.5Mb, 1Mb, 500kb, 250kb, 100kb, 50kb, 25kb, 10kb, and 5kb (Knight-Ruiz balancing by *Juicer tools*) resolutions. General genome organisation in the eight selected conditions was compared by plotting the coverage-corrected and the distance-and-coverage corrected Hi-C matrices.

#### Intra-chromosomal contact frequency analysis

We plotted the frequency of cis-chromosomal contacts in the raw data at various genomic distances. The frequency density was obtained by binning all cis-chromosomal contact distances (or, alternatively, the log10 distances), using 100 bins of equal size on a log_10_ genomic distance plot between 0 (1 bp) and 10 (10Gb).

#### Compartment analysis

The compartment signal was computed as the first principle component of the distance-and-coverage corrected interaction profile correlation matrix (Lieberman-Aiden *et al*, 2009) at 250kb resolution, with positive values aligned with H3K4me3 CHIP-seq on HeLa cells (https://www.encodeproject.org/experiments/ENCSR000DUA/ENCFF001XCR). For chromosomes for which the contact enrichment within chromosome arms was stronger than the compartment signal, we used the second principal component. The compartment signal for the selected conditions in each replicate was plotted for comparison, and the genome-wide concatenated ChIP-seq aligned principal components were clustered using hierarchical clustering (using 1 - Pearson correlation as the distance metric).

Using all A and B compartment bins defined by positive and negative compartment signal values in the control dataset, we also defined a compartmentalisation score, measuring the −log_2_ enrichment of trans-chromosomal contacts between all A and B compartment bins:

Compartment score = −log_2_ (2 *P*(A) *P*(B) *T* / *O*_AB_) where *T* = *O*_AA_ + *O*_AB_ + *O*_BB_ and *P*(A) = (2*O*_AA_ + *O*_AB_) / 2*T*, where *O*_AA_, *O*_AB_ and *O*_BB_ are the number of observed trans-chromosomal or long-cis (>2Mb) chromosomal contacts between all A-A, A-B and B-B bin pairs.

We defined 50 compartment groups of increasing A-compartment association, by binning all 250 kb bins of the genome into 50 equally sized groups according to their compartment signal in the G1-rep1 dataset. Group 1 contains 250kb bins with most negative compartment signal values (strongly B-compartment like) and group 50 contains 250kb bins with most positive compartment signal values (strongly A-compartment like). To obtain a pairwise contact enrichment matrix between these 50 groups, we computed the log_2_ enrichment of contacts between any two groups over the contact counts using a +10Mb-shifted compartment signal of the G1-rep1 dataset, with a correction to match the coverage profile of the 50 groups in the observed data.

#### Topological domain analysis

Topologically associating domains (TADs) were identified using HOMER v4.7. We computed directionality indices (Dis) (Dixon *et al*, 2012) of Hi-C interactions in 25kb sliding windows every 5kb steps, taking into account contacts to loci 1Mb upstream and downstream from the centre of the 25kb window, and smoothed the DI using a running average over a +/-25kb window. TADs were called between pairs of consecutive local maxima (start of a TAD) and minima (end of a TAD) of the smoothed DIs with a standard score difference (TAD ΔZ score) above 2.0, and the TAD ends were extended outward to the genomic bins with no directionality bias. HOMER calls TADs as well as TAD-less regions, so the genome coverage of the called TADs provides an estimate of domain organisation strength. For conditions with similar genome coverages, we could compare the numbers and sizes of TADs. To assess the average strength of TAD boundaries, we computed an average insulation score of the mean-shifted insulation score values in a 600kb window centered around those TAD boundaries.

#### Loop analyses

We identified loops genome-wide using the *HiCCUPS* algorithm of *Juicer tools* software (Durand *et al*, 2016). We called loops at 5kb, 10kb, and 25kb resolutions, employing Knight-Ruiz (KR) balancing and the default parameter values and a FDR threshold of 0.1, and merged these loop sets. Loops longer than 6Mb, caused by HeLa vs. hg19 assembly mapping artifacts such as translocations, were discarded. We note that the total number of loops depends on the number of replicates used, hence we only compared experiments with the same number of replicates.

We compared the total number of loops and their size distributions in the various conditions. We identified loop anchors by any genomic loci identified at a loop end with overlapping genomic coordinates, and calculated the frequency distribution of the number of loops from these loop anchors. Using the loop anchors and all loops connecting them, we defined connected networks (components, in graph theory) among all loops that are not connected by a loop to any outside genomic locus, and calculated the frequency distribution of loop numbers in these isolated networks of loops.

For measuring the aggregate peak enrichment of our loop sets, we used aggregate peak analysis. For loops in the 750kb-6Mb range, we used the *APA* algorithm of *Juicer tools* v0.7.5 to plot the sum of coverage-corrected Hi-C sub-matrices at 25kb resolution (Rao *et al*, 2014). The sub-matrices were centered and aligned around the peak coordinates of the looping loci such that the upstream loop anchor was at the center of the vertical, and the downstream loop anchor was at the center of the horizontal axis. The resulting plot displays a measure for the number of contacts that lie within the entire putative peak set at the center of the matrix, and the aggregation of contacts in a focal enrichments compared to the surroundings. The bottom left quarter of the plot displays the contact counts between loci between the looping anchors, characteristic of TAD-forming loops, and there is a general contact count decrease from the bottom left corner to the top right corner reflecting the distance dependent contact decay characteristic of the Brownian motion of linear polymers. To decouple this distance dependency, for loops that were exactly 300kb or 600kb long, we also plotted the average coverage-and-distance corrected Hi-C sub-matrices at 5kb resolution around the loops, using *HOMER*. Matrices were aligned as before, such that the diagonal of the sub-matrix also overlaps with the main diagonal of the Hi-C matrix. The plots show the contact enrichment compared to random contacts due to Brownian motion of the polymer. The selection of specific loop lengths enables more striking visualization of the domains bordered by some of the loops in the set. Another advantage of using the average rather than the sum of contacts is that it corrects for count differences due to different numbers of loops in the datasets, making the plots comparable between different loop sets.

To address the violation of the CTCF convergence rule in our datasets, we identified loops for which both anchors had consensus CTCF binding motifs overlapping with CTCF ChIP-seq peaks, as well as at least one SMC3 ChIP-seq peak. For these loops, we calculated the frequency of convergent, tandem and divergent CTCF sites in the control dataset, as well as for loops identified in WAPL, PDS5A/PDS5B and joint depleted datasets but not in control. For chains of loops, in which downstream loop anchors are also upstream loop anchors, we divided the loops into 5’, internal and 3’ loop categories (a 5’ loop’s upstream loop anchor only interacts downstream, and a 3’ loop’s downstream loop anchor only interacts upstream), and counted the frequency of forward and reverse CTCF binding motifs at these loop anchors.

## Acknowledgements

We thank Iain F Davidson and Benedikt Bauer for helpful discussions, Kristina Tabbada and VBCF for next generation sequencing, and Steven W. Wingett for Hi-C data mapping. Research in the laboratory of J.-M.P. is supported by Boehringer Ingelheim, the Austrian Science Fund (FWF special research program SFB F34 “Chromosome Dynamics” and Wittgenstein award Z196-B20), and the Austrian Research Promotion Agency (Headquarter grants FFG-834223). SS and PF are supported by the UK Biotechnology and Biological Sciences Research Council (BB/J004480/1), and CV and PF by the European Research Council Advanced Grant (DEVOCHROMO). KN is supported by an EMBO Long Term Fellowship (ALTF 1335-2016) and a HFSP fellowship (LT001527./2017). DAC acknowledges the VIPS Program of the Austrian Federal Ministry of Science and Research and the City of Vienna.

## Author contributions

GW, KN, DAC and JMP designed experiments and interpreted data together with CV, RS and SS. Cell lines were generated by MM and DAC (SCC1-GFP-AID with and without H2B-mCherry), WT (CTCF-GFP-AID) and NW, BK, MK (CAP-D2-mEGFP, CAP-D3-mEGFP). KN, DAC and GW performed RNAi, immunoblotting and microscopy experiments. GJ quantified chromatin compaction phenotypes. WT performed ChIP-seq experiments. MJH developed an algorithm for chromatin volume analysis. GW generated the Hi-C libraries with help from SS. CV and RS performed bioinformatic analyses of Hi-C data with input from GW. RS analyzed ChIP-seq data. JE, HZ, PF and JMP supervised the project. GW, CV, KN, DAC and JMP wrote the manuscript.

## Tables and their legends

See external Appendix Table 1 file.

**Appendix Table 1.**
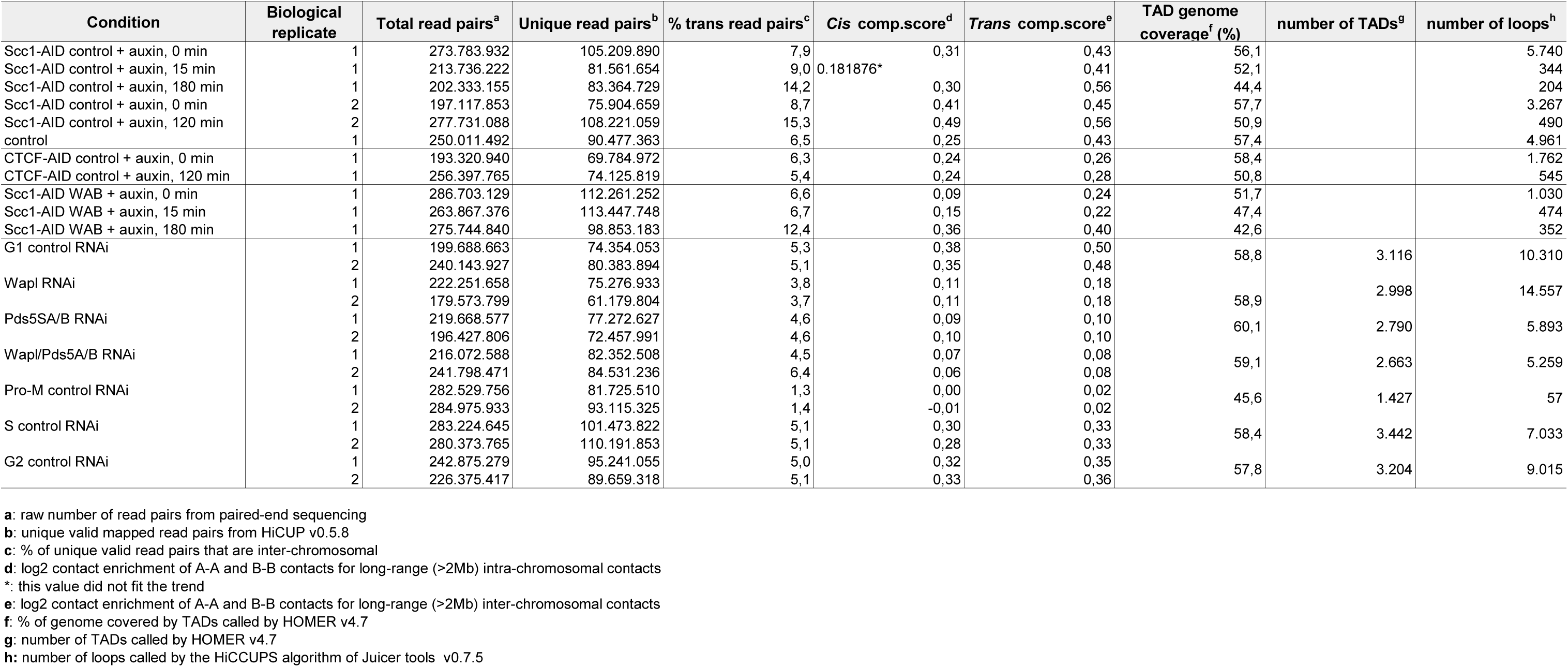
Summary statistics for Hi-C data sets generated in this study

## Expanded View Figure legends

**Figure EV1.**
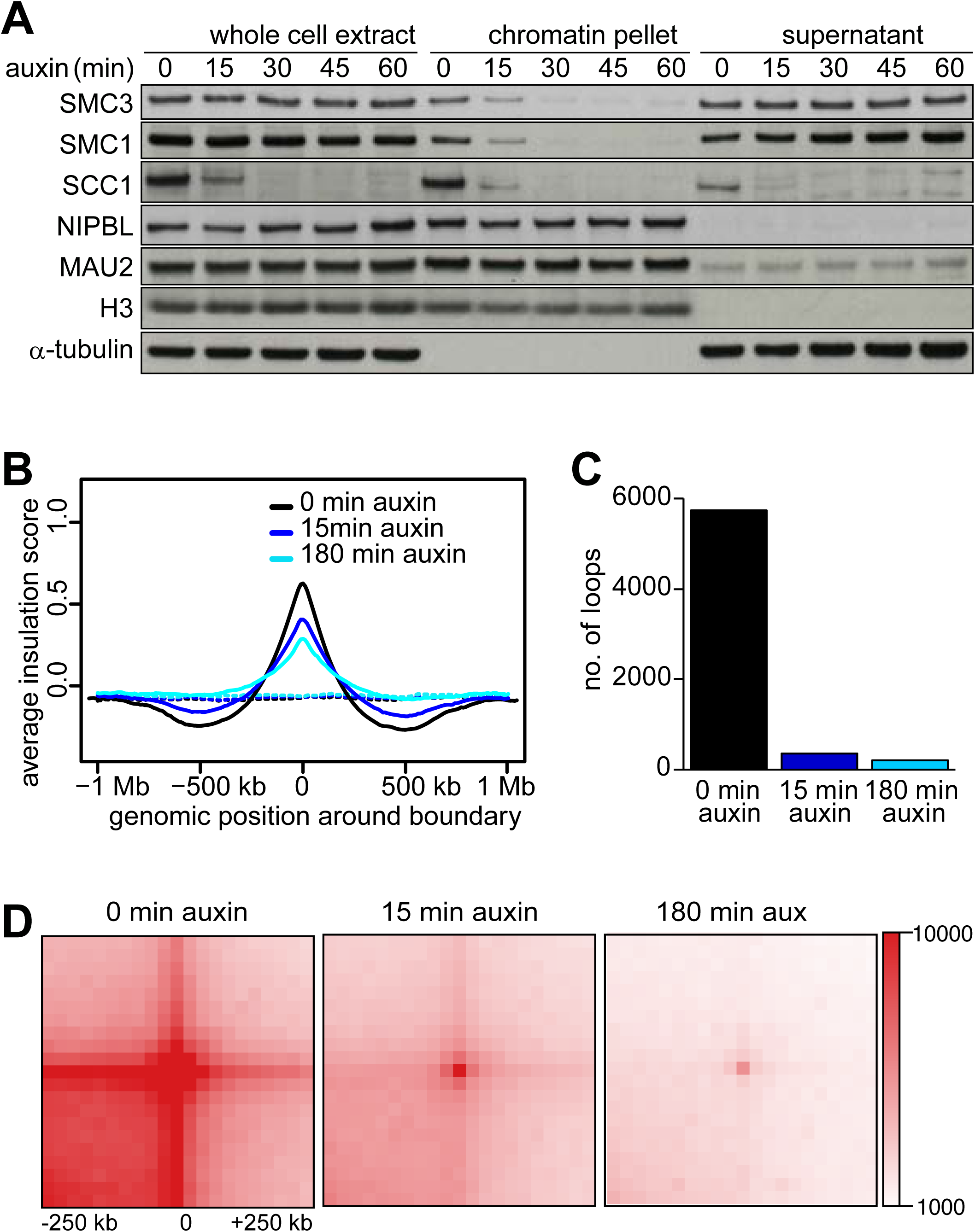
A. Same chromatin fractionation experiment as shown in Fig 1E. Note that the amount of cohesin loading complex, NIPBL and MAU2, on chromatin were largely unchanged after SCC1 degradation. B. Average insulation score around TAD boundaries identified for the SCC1-AID G1 control cells, at 0 (black), 15 (blue) and 180 minutes (cyan) after auxin addition. C. Number of loops identified by HICCUPS, at 0, 15 and 180 minutes after auxin addition after auxin addition. Colours are the same as in (B). D. Total contact counts around loops after auxin addition, for all 750kb-6Mb long loops identified by HICCUPS in G1 control.

## Appendix Figure legends

**Appendix Figure S1.**
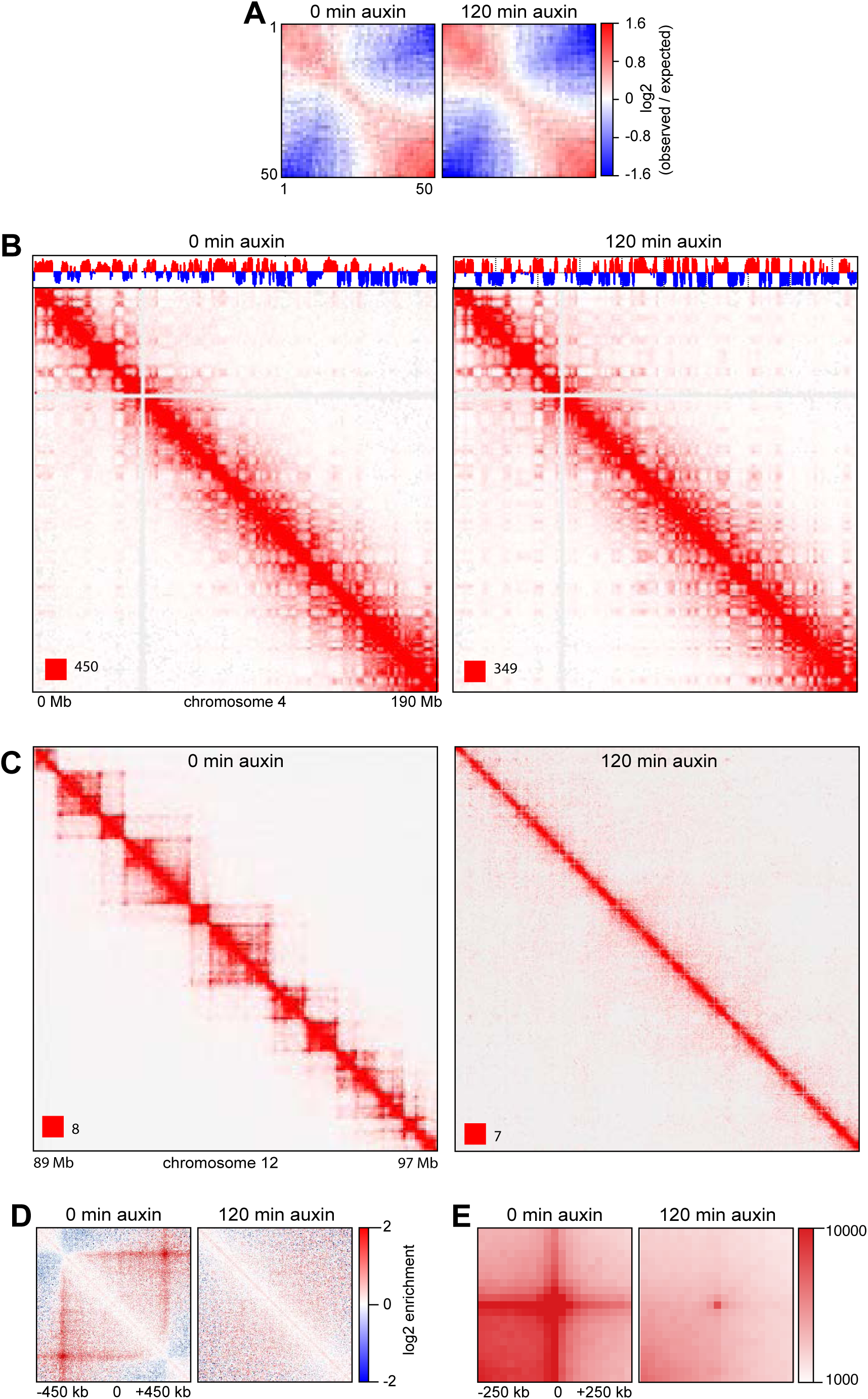
A. Inter-chromosomal contact enrichment between 250kb bins with varying compartment strength from most B-like (1) to most A-like (50), in SCC1-AID control cells in a replicate experiment, at 0 and 120 minutes after auxin addition. B. Coverage-corrected Hi-C contact matrices of chromosome 4, at 0 and 120 minutes after auxin addition in SCC1-GFP-AID cells. The corresponding compartment signal tracks at 250kb bin resolution are shown above the matrices. The matrices were plotted using Juicebox. C. For the same conditions, coverage-corrected Hi-C contact matrices in the 89-97Mb region of chromosome 12, plotted by using Juicebox. D. Average contact enrichment around loops after auxin addition in SCC1-GFP-AID cells, for the 82 x 600 kb long loops identified by HICCUPS in G1 control. The matrices are centered (0) around the halfway point of the loop anchor coordinates. E. Total contact counts around loops after auxin addition in SCC1-GFP-AID cells, for all 750kb-6Mb long loops identified by HICCUPS in G1 control.

**Appendix Figure S2.**
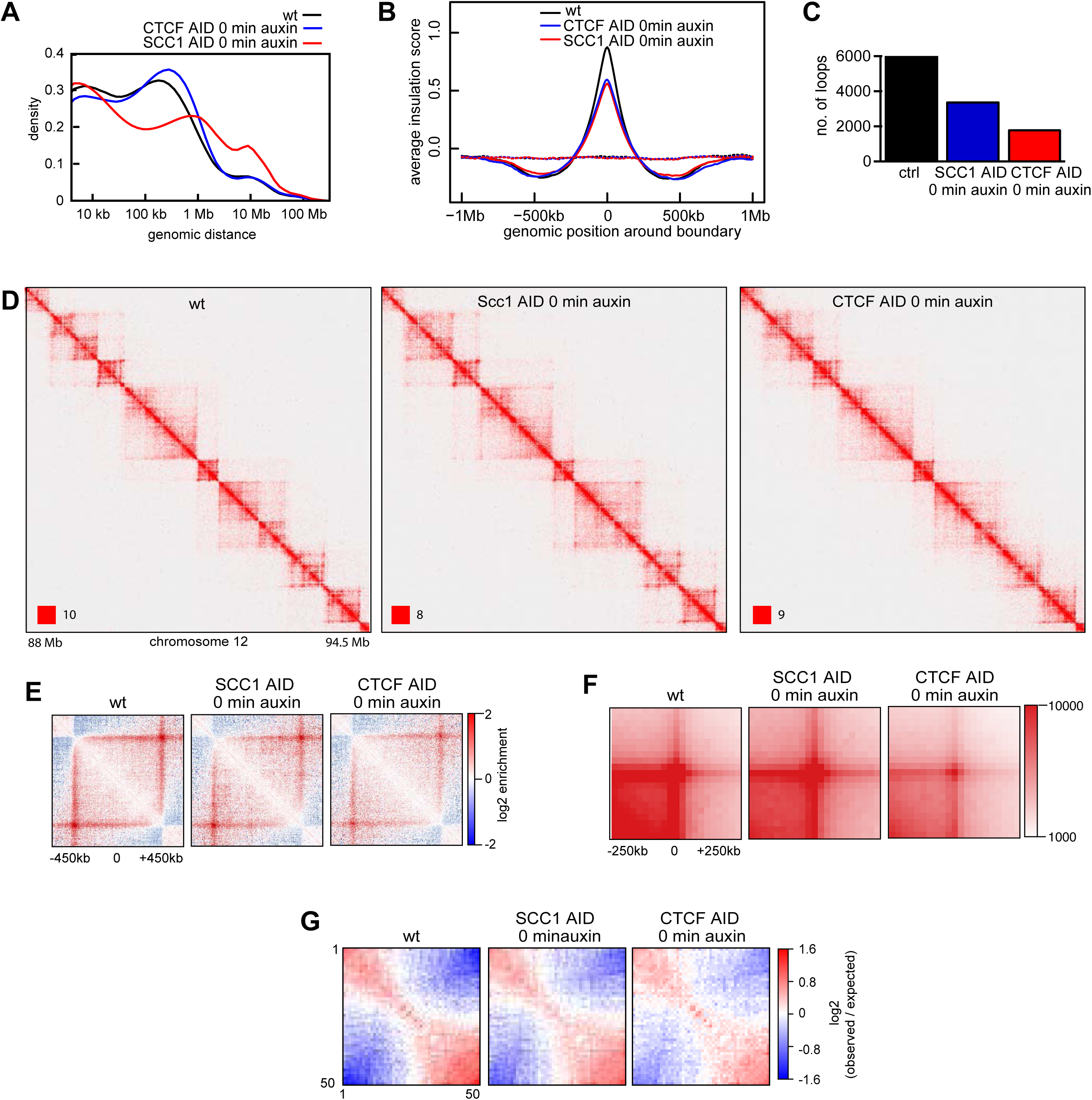
A. Intra-chromosomal contact frequency distribution as a function of genomic distance, in the wild type (WT, black), CTCF-AID (red) and SCC1-AID (blue) cells immediately after auxin addition. B. Average insulation score around TAD boundaries identified in G1 control cells, for the same conditions as in (A). C. Number of loops identified by HICCUPS, for the same conditions. D. Coverage-corrected Hi-C contact count matrices in the 88-94.5Mb region of chromosome 12, for the same conditions. E. Average contact enrichment around loops after auxin addition, for the 82 x 600 kb long loops identified by HICCUPS in G1 control. F. Total contact counts around loops after auxin addition, for all 750kb-6Mb long loops identified by HICCUPS in G1 control. G. Inter-contact enrichment between bins with varying compartment strength from most B-like (1) to most A-like (50).

**Appendix Figure S3.**
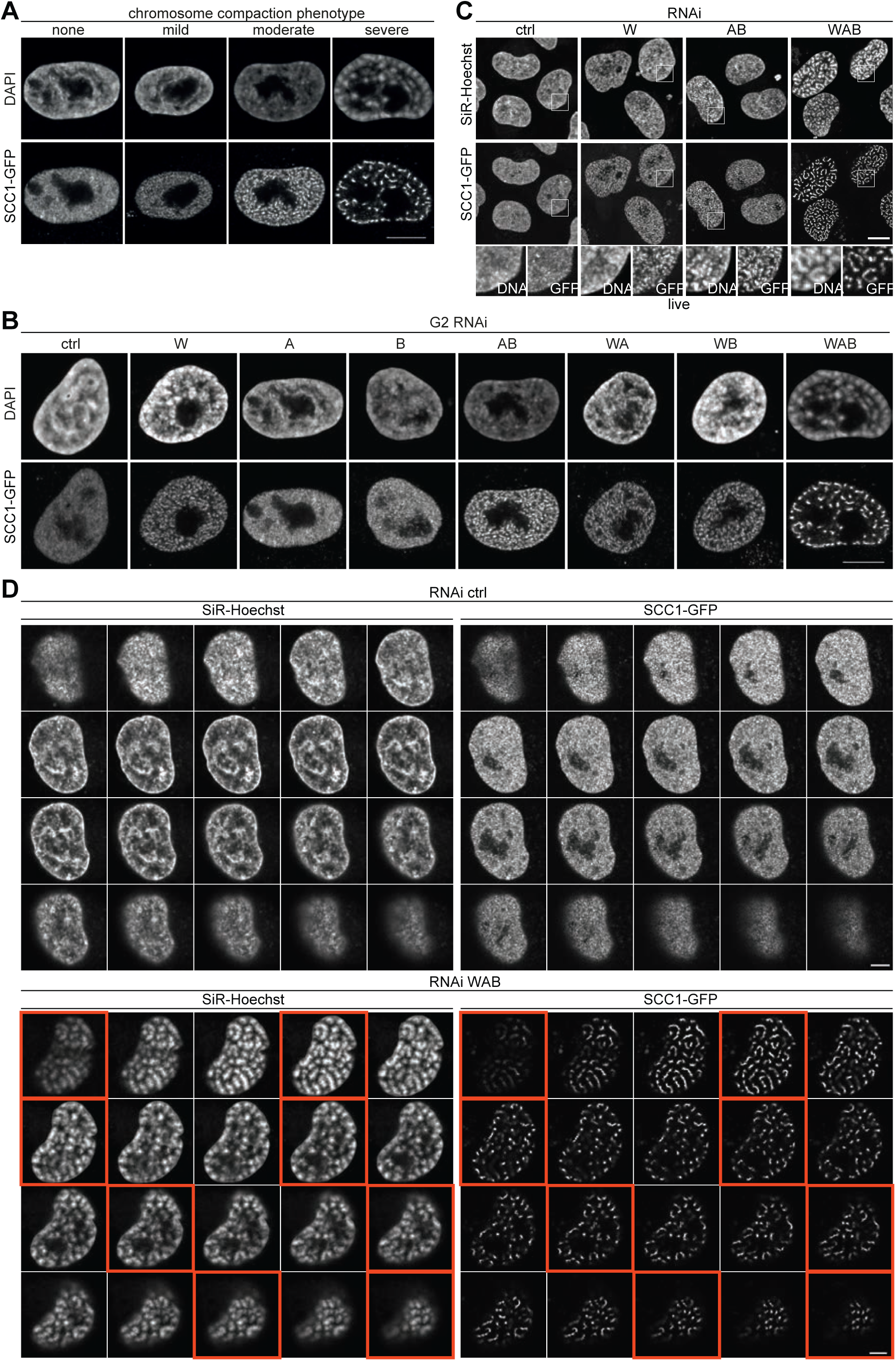
A. Representative images used for phenotypic classification analysis shown in Fig 3C. Scale bar shows 10 μm. B. Representative immunofluorescence images of SCC1-GFP cells in G2 phase stained for GFP. DNA was counterstained with DAPI. Scale bar shows 10 μm. C. Live cell imaging of SCC1-GFP cells that had been depleted for control, W, AB and WAB. DNA was stained by SiR-DNA by Spirochrome. Three of the confocal sections are shown by perspective view. Scale bar shows 10 μm. D. Live cell imaging of SCC1-GFP cells that were depleted for WAB or control depleted. Individual confocal sections are shown (confocal distance between original sections = 0.4 μm). The images entangled by the orange lines are also shown in Fig 3E. DNA was counter stained with SiR-DNA by Spirochrome. Scale bar shows 5 μm.

**Appendix Figure S4.**
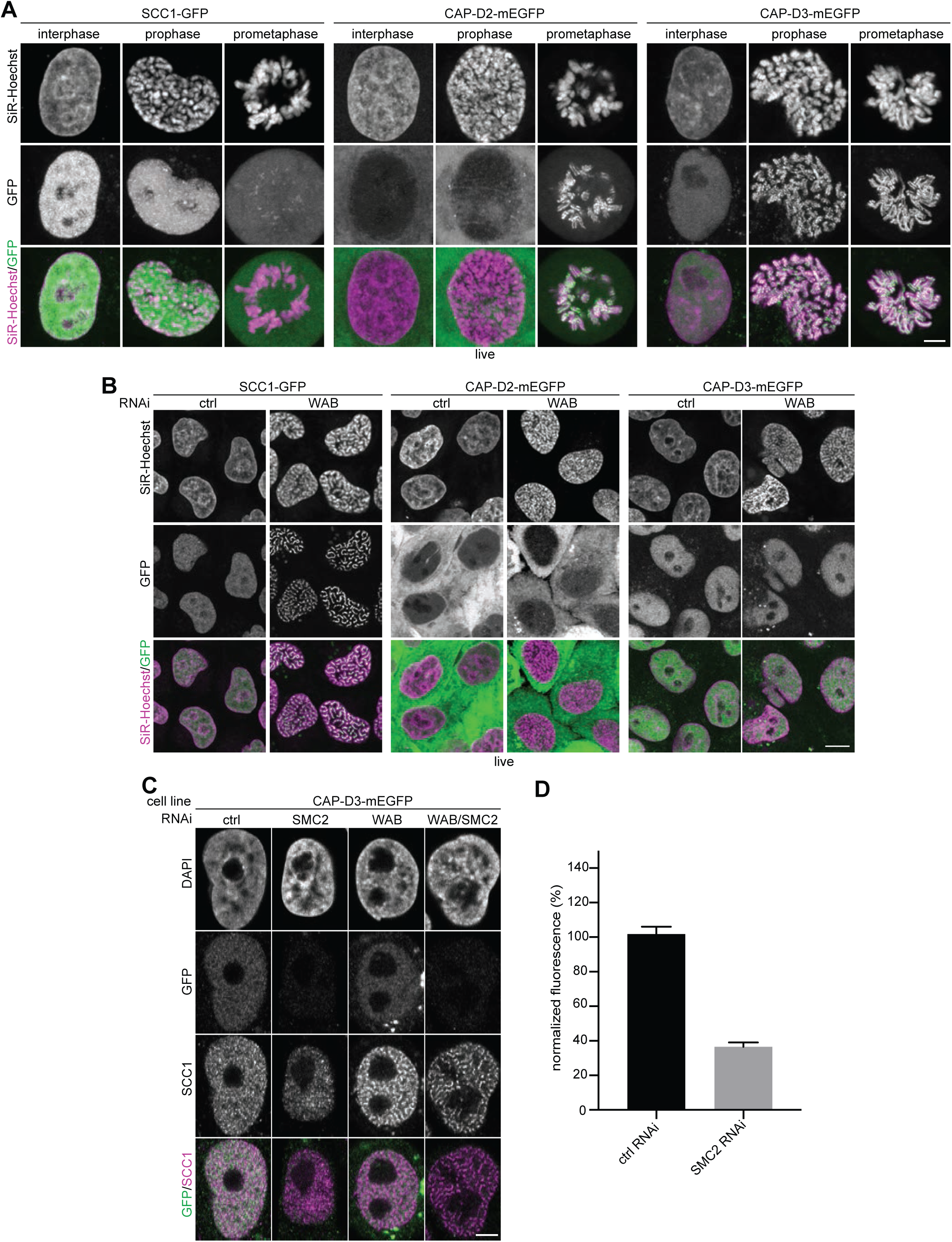
A. Live cell imaging of SCC1-GFP, CAP-D2-mEGFP and CAP-D3-mEGFP cells in which all alleles of these genes are C-terminally fused to a GFP tag by CRISPR/Cas9 engineered genome editing. Representative cells at interphase, prophase and prometaphase are shown. DNA was stained with SiR-DNA by Spirochrome. Scale bar indicates 5 μm. B. Live cell imaging of SCC1-GFP, CAP-D2-mEGFP and CAP-D3-mEGFP cells depleted for control or WAB by RNAi. DNA was stained by SiR-DNA by Spirochrome. Scale bar indicates 10 μm. C. Immunofluorescence images of CAP-D3-mEGFP cells depleted for SMC2, WAB or SMC2/WAB. GFP staining was used as readout for estimating condensin depletion efficiency. Scale bar indicates 5 μm. D. Quantification of immunofluorescence intensities for GFP signals in cells depleted for control and SMC2 is shown. Error bar depicts standard deviations of the means.

**Appendix Figure S5.**
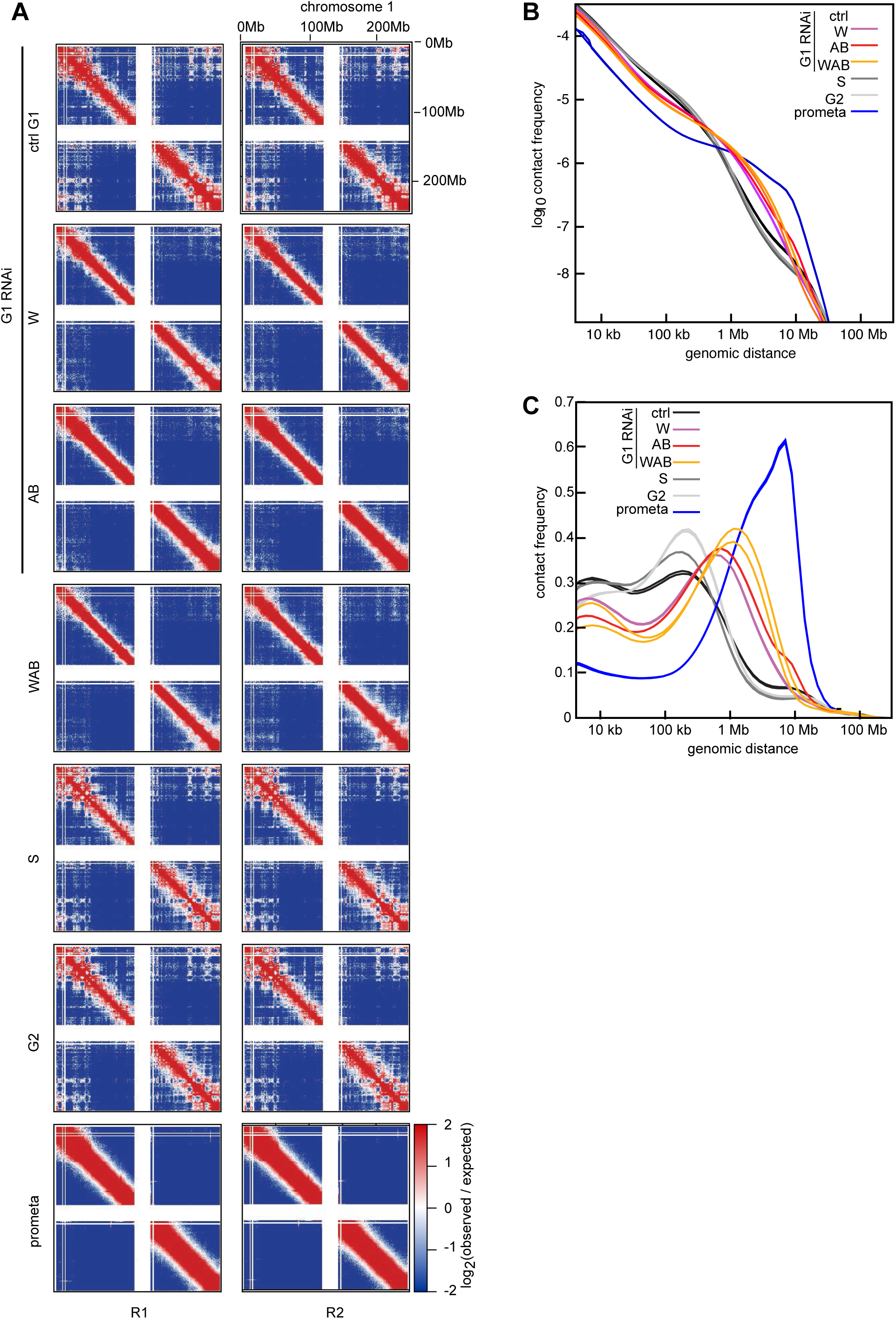
A. Coverage-corrected Hi-C contact enrichment matrices (using HOMER) of chromosome 1, for the same conditions as in Fig 5, showing the two biological replicates side by side. B-C. Intra-chromosomal contact frequency distribution as a function of genomic distance using equal sized (B) or logarithmically increasing (C) genomic distance bins, for the two biological replicates. Colours are the same as in Fig 5.

**Appendix Figure S6.**
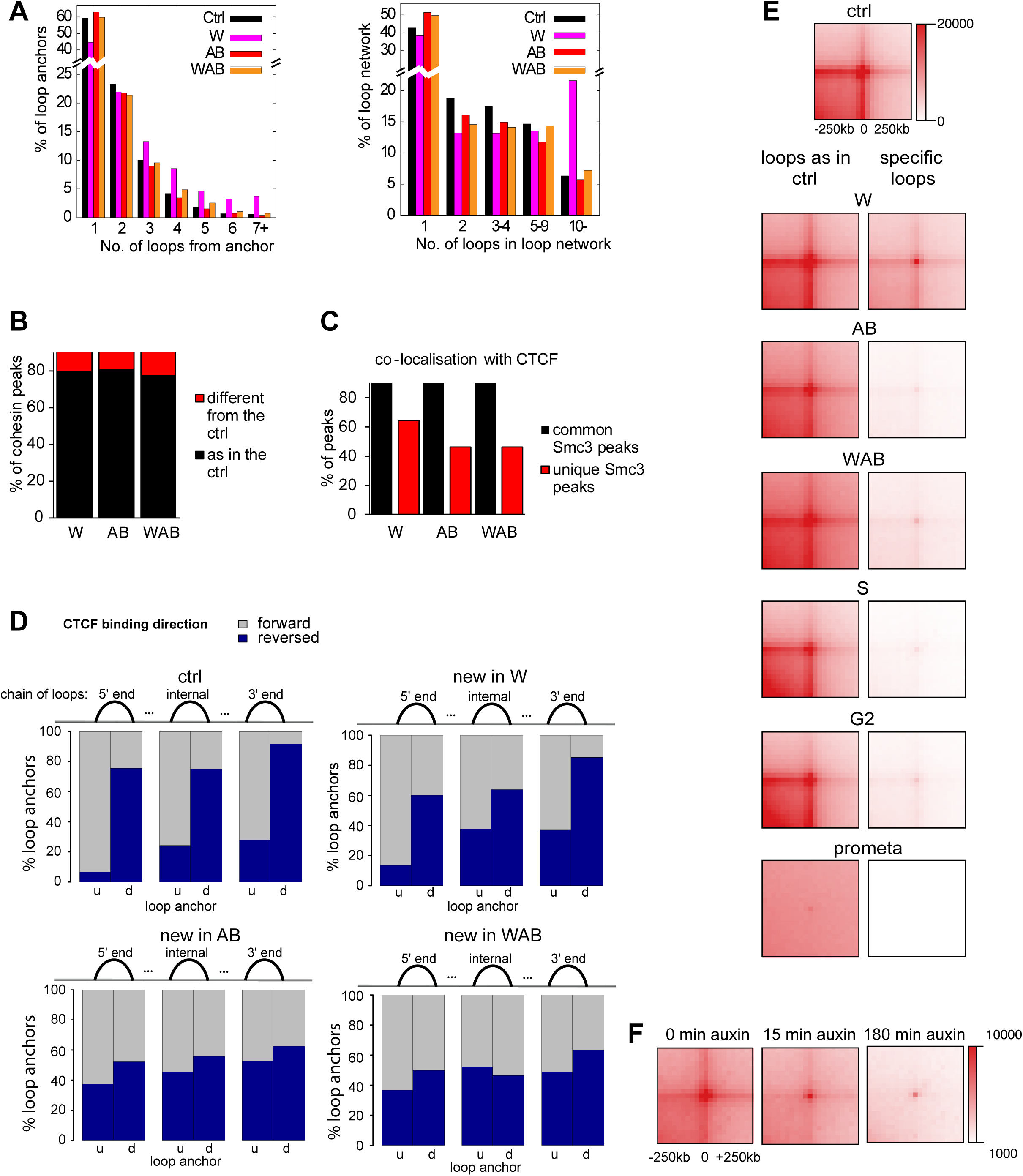
A. The frequency distributions of the number of loops formed from all loop anchors (left), and the number of loops forming all contiguous loop networks (right), for the G1 control (ctrl, black), WAPL depleted (W, magenta), PDS5A/B depleted (AB, red) and WAPL/PDS5A/B depleted (WAB, orange) RNAi samples. B. Percentages of cohesin peaks in each of the three RNAi depleted conditions which are either common with G1 control (black) or newly arising in the particular depleted condition (red). C. Comparison of cohesin sites in the three RNAi depleted conditions regarding their co-localization with CTCF. Black bars indicate the percentage of overlap with CTCF for those cohesin sites which are common between the particular condition and G1 control. Red bars show the overlap with CTCF for cohesin sites which arise newly in the particular condition and are not found in control. D. For 5’, internal and 3’ loops of loop networks, the proportion of their upstream (u) and downstream (d) loop anchors with forward (grey) and reverse (blue) CTCF binding orientation. The same loops as in Appendix Fig S6F were used. E. Total contact counts around loops for all 750kb-6Mb long loops identified by HICCUPS in G1 control cells (left), or in the corresponding sample but not in G1 control cells (right). F. Total contact counts around loops after auxin addition in the WAPL/PDS5A/B RNAi depleted SCC1-AID cell line, 0, 15 and 180 minutes after auxin addition, for all 750kb-6Mb long loops identified by HICCUPS in G1 control.

